# AAV2/9-mediated gene transfer into murine lacrimal gland leads to a long-term targeted tear film modification

**DOI:** 10.1101/2022.02.16.480632

**Authors:** Benoit Gautier, Lena Meneux, Nadège Feret, Christine Audrain, Laetitia Hudecek, Alison Kuony, Audrey Bourdon, Caroline Le Guiner, Véronique Blouin, Cécile Delettre, Frédéric Michon

## Abstract

Corneal blindness is the fourth leading cause of blindness worldwide. Since the corneal epithelium is constantly renewed, non-integrative gene transfer cannot be used to treat corneal diseases. In numerous of these diseases, the tear film has been reported to be defective. Tears are a complex biological fluid secreted by the lacrimal apparatus. Their composition is modulated according to the context. For instance, after a corneal wound, the lacrimal gland secretes reflex tears, which contain specific growth factors supporting the wound healing process. In specific pathological contexts, such as dry eye diseases, the tear composition can support neither corneal homeostasis, nor wound healing. Here, we propose to use the lacrimal gland as bioreactor to produce and secrete specific factors to support corneal physiology. In this study, we used an AAV2/9-mediated gene transfer to supplement the tear film. First, we demonstrate that a single injection of AAV2/9 is sufficient to transduce all epithelial cell types of the lacrimal gland efficiently and widely. Then, we show that lacrimal gland physiology and corneal integrity are maintained after the injection of an AAV2/9-mediated nerve growth factor expression in the lacrimal gland. Remarkably, this injection induces an important and long-lasting secretion of this growth factor in the tear film. Altogether, our findings provide a new clinically applicable approach to tackle corneal blindness.

## Introduction

To ensure clear vision, the anterior parts of the eye, namely the cornea and lens, need to be fully transparent. While the lens is protected by being located inside the eye, the cornea is the most external tissue of the eye, and thus prone to environmental aggressions. This ectodermal organ is subjected to life-long cell renewal, which relies on stem and progenitor cells^1^. To coordinate epithelium homeostasis, corneal microenvironment is composed of epithelial cell-cell communication, dense innervation, and tear film. The latter is the source of corneal hydration and nutrients for the epithelium^2^. Moreover, after wounding, the tear composition changes to support corneal wound healing, through a modification of the factors secreted by the lacrimal gland (LG)^3^.

The tear composition change requires an efficient sensory network of the cornea. The dense corneal innervation is essential to maintain corneal physiology. First, blinking and tear composition adaptation depends on the activity of corneal sensory nerves. Furthermore, the neurotrophic factors released by the nerves for the epithelium are crucial for homeostasis and wound healing^4^. Among the most prominent causes for corneal defects, three are making most of the influx of patients in hospital. First cause, physical wounds, such as abrasions^5^, are mostly caused by small foreign objects that scratch the epithelium. This injury is painful, and the subsequent oedema provokes photophobia and impairs visual acuity. If the particle gets embedded within the epithelium, corneal irregularities might form, resulting in continuous pain and significant visual disruption^6^. Dry eye diseases account for the second cause of corneal defects. Dry eye diseases can arise from genetic disease, such as Gougerot-Sjörgren syndrome^7^, or from ageing, as up to 1/3 of the elder population can be affected^8, 9^. The altered tear film has an imbalanced composition, offering less nutrients and growth factors to the corneal epithelium, which in turn affects corneal homeostasis. Consequently, persistent epithelial defects appear, such as ulceration, melting and perforation, impairing sight^10^. Third cause, neurotrophic keratitis is due to a partial or complete loss of corneal innervation, causing a defected corneal homeostasis^11^. Neurotrophic keratitis results from defected corneal wound healing after abrasion or transplant, from neurodegenerative diseases, or from chronic metabolic diseases, such as diabetes. Currently 415 million adults globally are diagnosed with diabetes, and the World Health Organization projected that there will be 640 million adults by 2040^12^. While being underdiagnosed, diabetic keratopathy affects 47 to 64% of diabetic adults^13^. The main symptom of neurotrophic keratitis is corneal ulceration and perforation.

The current treatments for these corneal defects are topical and consist in eye drops, which can be supplemented with autologous serum^14^, or nerve growth factor (NGF) in the case of neurotrophic keratitis^15^. Not only these treatments are heavy for patients, especially autologous serum, but the frequent lack of patient compliance to a prescribed eyedrop regimen lead to increased sight defects^16^.

These treatments solely rely on the addition of an external eye drop solution mimicking the tear film composition without considering the LG which produces, secretes, and modulates the tear film composition. Consequently, using LG directly as a bioreactor would constitute an appealing alternative strategy to modulate the tear film composition. This could be achieved by adenovirus-associated virus (AAV) vector-mediated gene transfer into LG. Indeed, AAV vectors present many advantages for gene delivery. They efficiently transduce a broad range of cells in which they allow long-lasting transgene expression. Importantly, they trigger limited/mild immunogenic responses *in vivo* which overall asserts their biosafety^17^. Numerous AAV-based gene therapies have emerged as evidenced by many ongoing clinical trials, namely for neurodegenerative, neuromuscular, cardiovascular, ocular genetic diseases and cancer^18, 19^. Only three AAV-based gene therapies have been approved by the FDA, among which two are still used. Luxturna® (voretigene neparvovec) is used for the treatment of biallelic RPE65 mutation-associated retinal dystrophy^20^ while Zolgensma® (onasemnogene abeparvovec-xioi) is delivered to paediatric patients under two years of age suffering from spinal muscular atrophy^21^. Importantly, in the field of ophthalmology, the majority of studies using AAV-based gene delivery have focussed on retina and to a lesser extent on cornea, while very few data are available on AAV-based gene transfer into LG^22–24^.

In this study, we present a novel strategy to modulate corneal physiology through targeted tear film modification by transferring a gene of interest into LG. First, we confirmed that AAV-mediated gene transfer was feasible in murine LG using AAV2/5 or AAV2/9. After demonstrating that all epithelial cell types are prone to AAV-mediated gene transfer, we used AAV2/5 and AAV2/9-mediated murine nerve growth factor (mNGF), as proof of concept, to establish the parameters for efficient gene transfer, allowing a targeted modification of tear composition. We investigated the impact of AAV serotype impact on secreted protein level and chose AAV2/9 vector to investigate the duration of tear film modulation, as well as the safety of such an approach on the cornea. Altogether, our results demonstrate that a single AAV2/9 injection into murine LG could be used to specifically modify the tear film and consequently to support corneal physiology in pathological contexts, such as neurotrophic keratitis, or recurring abscesses.

## Results

### AAV2/9 and AAV2/5 mediate an efficient gene transfer in the lacrimal gland

To investigate the use of the LG as bioreactor for protein secretion in the tear film, we first established an injection protocol that allows an efficient AAV-mediated gene transfer after a single injection into murine LG. We used AAV2/9-CAG-GFP and AAV2/5-CAG-GFP to monitor the extent of vector diffusion within LG. The immunostaining of GFP demonstrated that these AAV serotypes induced GFP expression in all territories of the LG (Figure 1). To rule-out a possible tropism of AAV serotypes for a specific-cell type, we looked at GFP expression, one month after injection, in two different LG epithelial cell populations, namely the E-Cadherin (E-Cadh) positive cells, located in the acinar compartment, and the Keratin19 (Krt19) positive subpopulation, specifically localized in the ducts^25^. The GFP / E-Cadh co-labelling showed that almost all acinar cells were GFP positive regardless of the used serotype (Figure 2A, S1). Similarly, the GFP / Krt19 co-labelling demonstrated that most of the ductal cells were GFP positive after AAV2/9-CAG-GFP and AAV2/5- CAG-GFP injection (Figure 2B, S2).

**Figure 1.**
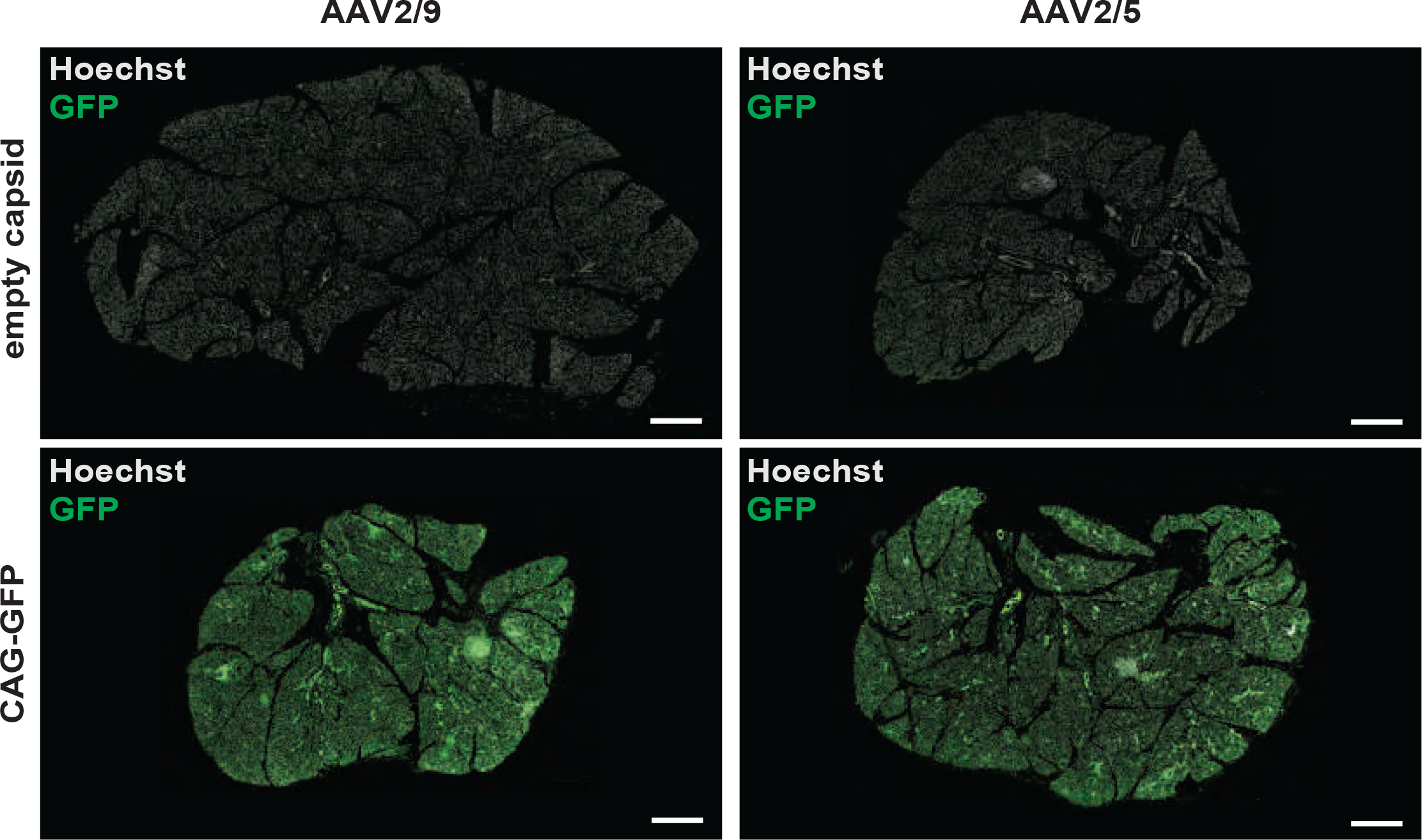
AAV2/9 and AAV2/5-CAG-GFP transduce widely the murine LG. Representative images of LG longitudinal sections show GFP protein (green) following a single injection of AAV2/9 or AAV2/5-CAG-GFP into murine LG, when compared to AAV2/9 or AAV2/5 empty capsid-injected mice respectively (10^10^ vg / LG in 3 µl, *n* = 3 mice per group). Mice were sacrificed one month post injection. Nuclei are counterstained with Hoechst 33342 (white). Scale bar: 500 µm.

**Figure 2.**
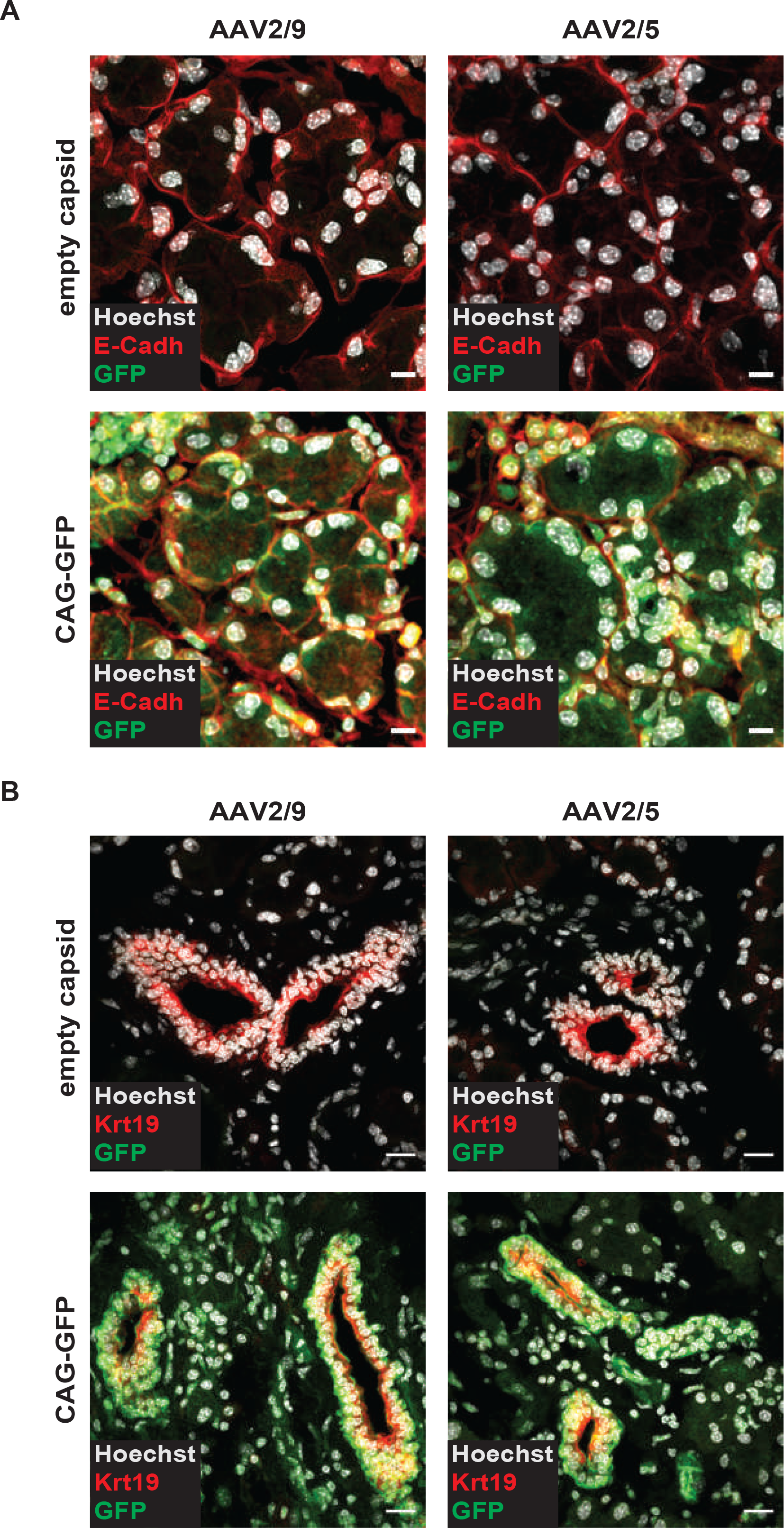
AAV2/9 and AAV2/5-CAG-GFP transduce LG acinar and ductal cells. Representative images of LG longitudinal sections show GFP protein (green) in acinar cells immunostained for E- Cadh (red, **A**) and in ductal cells immunostained for Krt19 (red, **B**) after a single injection of AAV2/9 or AAV2/5-CAG-GFP into murine LG, when compared to AAV2/9 or AAV2/5 empty capsid- injected mice respectively (10^10^ vg / LG in 3 µl, *n* = 3 mice per group). Mice were sacrificed one month post injection. Nuclei are counterstained with Hoechst 33342 (white). Scale bar: 10 µm (**A**) and 20 µm (**B**).

To evaluate a possible discrepancy between AAV serotypes 2/5 and 2/9 in the transduction efficiency, we analyzed by Western-blot the levels of GFP protein expressed in LG after injection of AAV2/9- or AAV2/5-CAG-GFP (Figure 3). As expected, while in the LG injected with AAV2/9 or AAV2/5 empty capsids, no GFP protein was detected (Figure 3A), whereas high levels of GFP were detected in the LG injected with AAV2/9 or AAV2/5-CAG-GFP respectively. Interestingly, when comparing the two AAV serotypes, GFP levels were found significantly higher after AAV2/9- CAG-GFP injection than after AAV2/5-CAG-GFP injection (Figure 3B).

**Figure 3.**
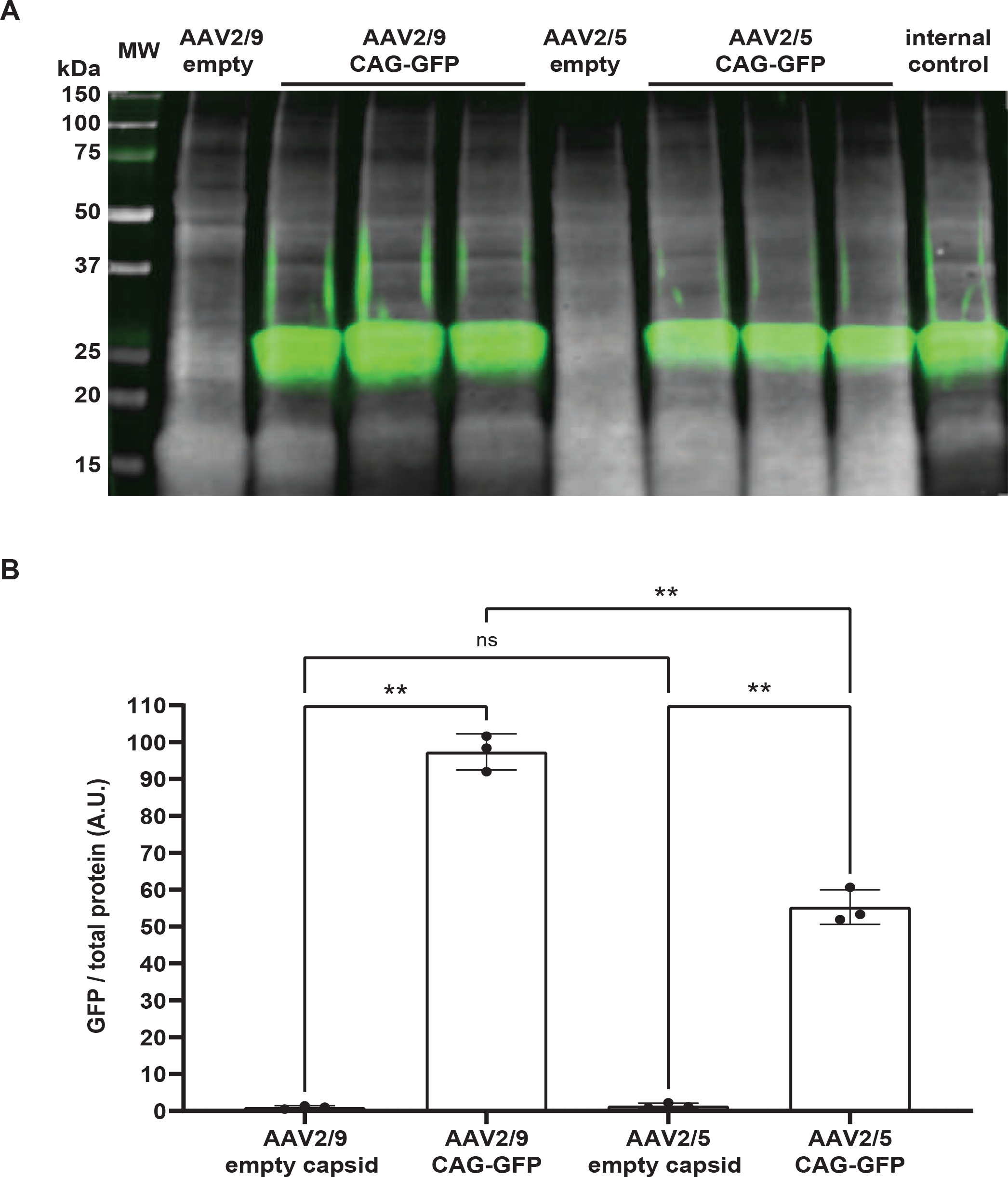
GFP is detected in AAV2/9 and AAV2/5-CAG-GFP-injected LG and dosed by Western blot. **A.** Representative Western blot image showing GFP level (green) and total protein as loading control (white) in murine LG lysates from AAV2/9 or AAV2/5-CAG-GFP-injected mice, when compared to AAV2/9 or AAV2/5 empty capsid-injected mice respectively (10^10^ vg / LG in 3 µl). Mice were sacrificed one month post injection. **B.** Quantification of GFP / total protein ratio in LG lysates (*n* = 3 mice per group). Results are expressed as the mean ± SD. Statistical analysis using Brown-Forsythe and Welch ANOVA tests followed by Dunnett’s T3 multiple comparisons test. ** *p* = 0.0026 between AAV2/9 empty capsid and AAV2/9 CAG-GFP groups, ** *p* = 0.0077 between AAV2/5 empty capsid and AAV2/5-CAG-GFP groups, ** *p* = 0.0018 between AAV2/9- CAG-GFP and AAV2/5-CAG-GFP groups; ns, not significant; A. U., arbitrary units. Source data are provided as a Source Data file.

Taken together, our results demonstrate that LG can be widely transduced by AAV2/9 and AAV2/5, and that serotype 2/9 exhibits a higher transduction efficiency.

### The transgene expression leads to the secretion of the resulting protein in the tear film

To evaluate if AAV2/9 and AAV2/5 serotypes have a different impact on the amount of protein secreted in the tear film, we chose to express the murine nerve growth factor (mNGF) in the LG, using either AAV2/9 (AAV2/9-CAG-mNGF) or AAV2/5 (AAV2/5-CAG-mNGF). Following AAV injection into LG, the transgene expression should lead to the secretion of mNGF in the tear film. To assess this secretion, we measured mNGF level in the tear film, using ELISA and Western-blot analysis (Figure 4). Tear film contains a basal level of mNGF^26^, which can have two forms^27, 28^. The pro-mNGF, of higher molecular weight is cleaved to generate the mature mNGF, the active form. As expected from the GFP results, injection of AAV2/9 and AAV2/5 CAG-mNGF induced a significant increase of the mNGF levels in the tear film, whereas only endogeneous basal expression levels of mNGF were detected with the empty AAV vectors. Moreover, consistant with the GFP expression data in LG, mNGF amount in the tear film, was more than 3-fold higher after AAV2/9 injection than after AAV2/5 injection (Figure 4). Moreover, Western blot analysis confirmed that injection of AAV2/9 and AAV2/5 CAG-mNGF led to a significant increase of the total mNGF level in the tear film (Figure 5A and 5B). Importantly, this analysis revealed that mNGF was consistently found in the tear film in its pro-mNGF form (Figure 5C). However, the analysis of the mature mNGF showed a maturation process (Figure 5D). Indeed, the mature mNGF form was close to absent from the tear film of mice injected in LG with empty AAV2/9 or AAV2/5, whereas a large amount of mature mNGF was detected after AAV-mNGF gene transfer. Notably, AAV2/9- CAG-mNGF gave rise to 4-times more of the mature form than with AAV2/5-CAG-mNGF injection (Figure 5D).

**Figure 4.**
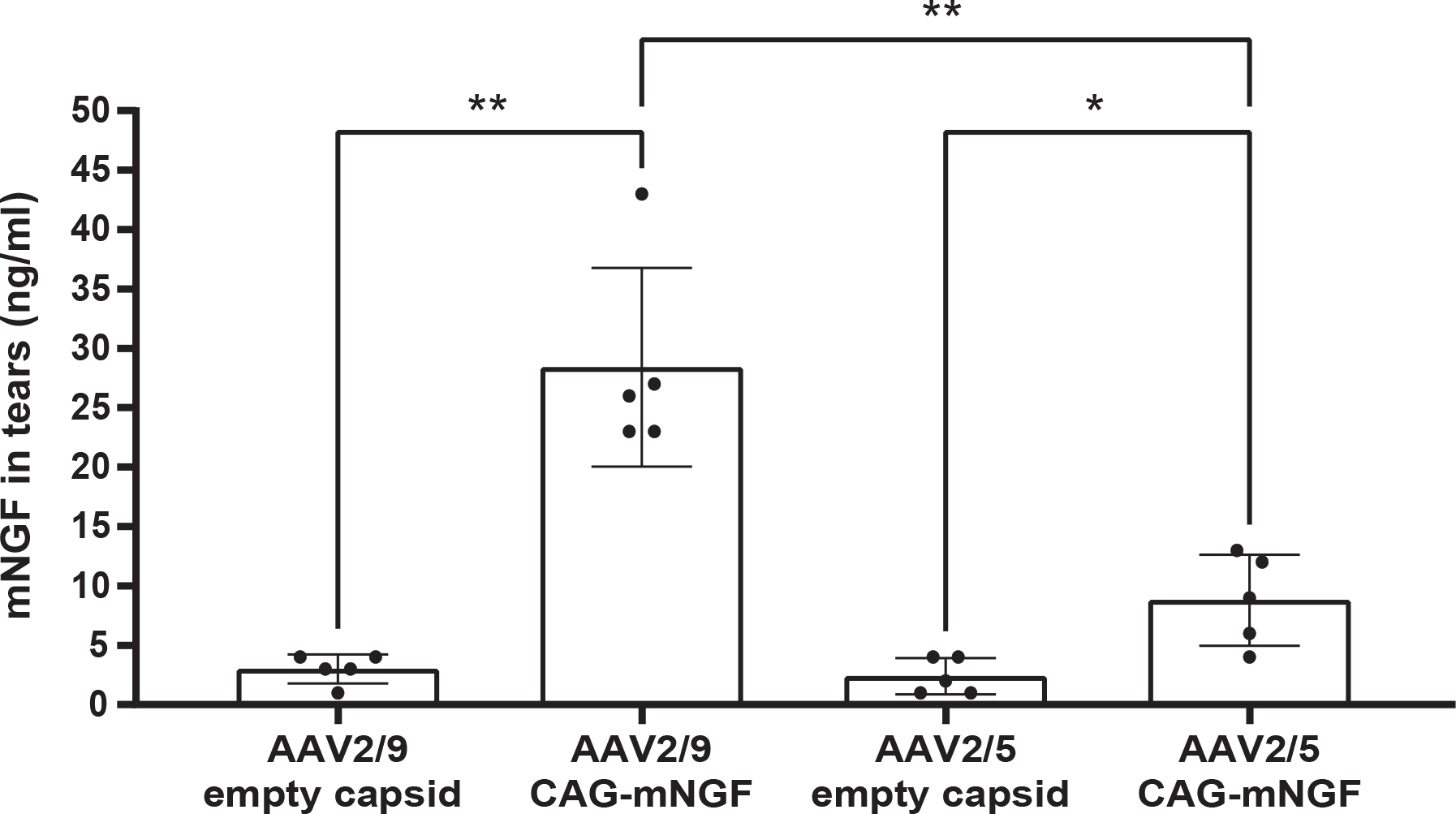
Injections of AAV2/9 and AAV2/5-CAG-mNGF into murine LG significantly increase the level of mNGF in tears. mNGF protein levels (ng / ml) were measured by ELISA in the tears of AAV2/9 or AAV2/5-CAG-mNGF-injected mice, when compared to AAV2/9 or AAV2/5 empty capsid-injected mice, one month post injection respectively (10^10^ vg / LG in 3 µl, *n* = 5 mice per group). Results are expressed as the mean ± SD. Statistical analysis using Brown-Forsythe and Welch ANOVA tests followed by Dunnett’s T3 multiple comparisons test. ** *p* = 0.0065 between AAV2/9 empty capsid and AAV2/9-CAG-mNGF groups, * *p* = 0.0464 between AAV2/5 empty capsid and AAV2/5-CAG-mNGF groups, ** *p* = 0.0085 between AAV2/9-CAG-mNGF and AAV2/5- CAG-mNGF groups. Source data are provided as a Source Data file.

**Figure 5.**
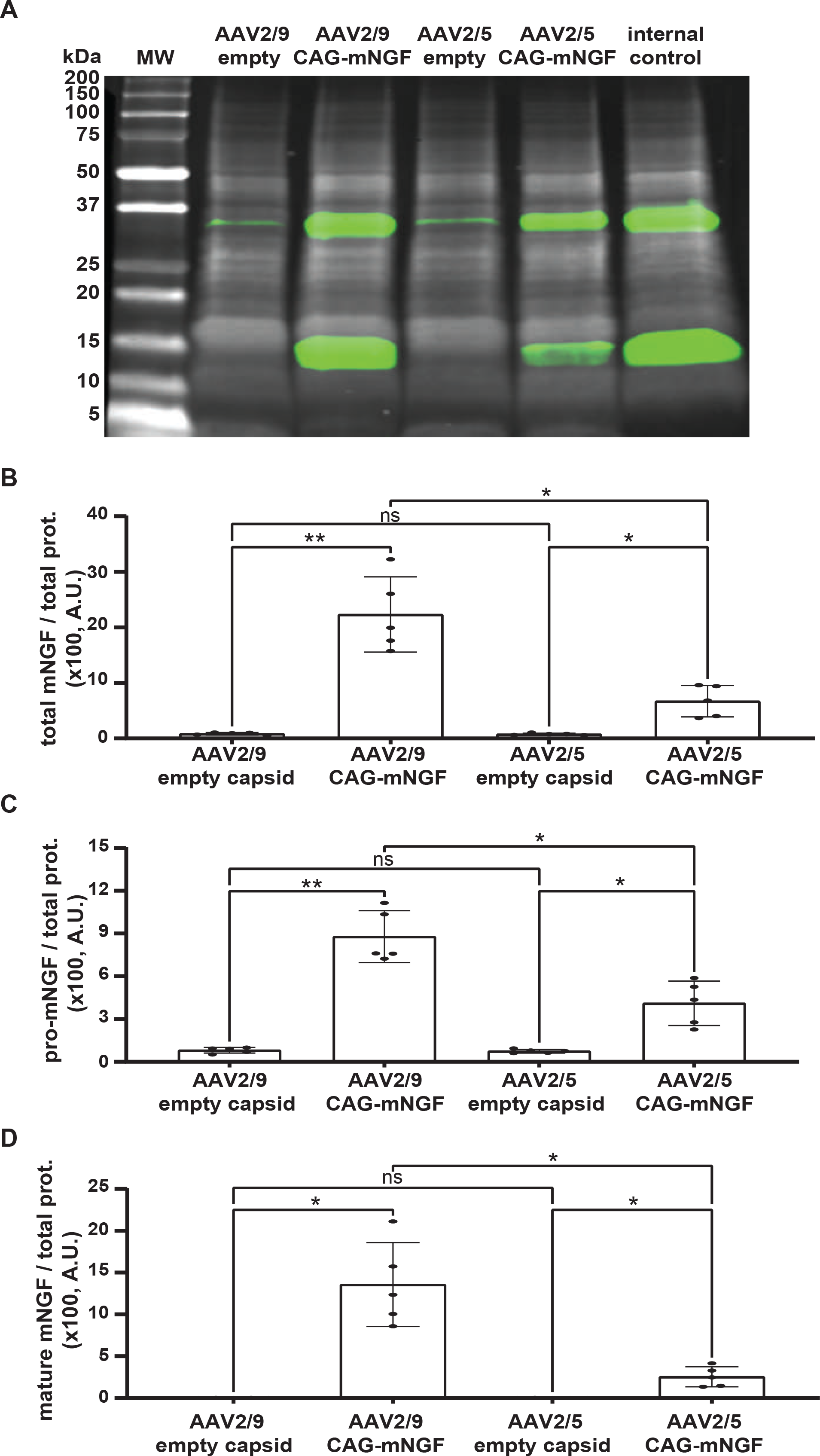
Injections of AAV2/9 and AAV2/5-CAG-mNGF into murine LG significantly increase the level of total, pro and mature mNGF in tears. mNGF protein levels were analyzed by Western blot experiments in the tears of AAV2/9 or AAV2/5-CAG-mNGF-injected mice, when compared to AAV2/9 or AAV2/5 empty capsid-injected mice, one month post injection respectively (10^10^ vg / LG in 3 µl, *n* = 5 mice per group). **A**. Representative Western blot image showing mNGF level (green) and total protein as loading control (white) in the tears of injected mice. Quantification of total mNGF (**B**), pro mNGF (**C**) and mature mNGF (**D**) protein levels in the tears of injected mice Results are expressed as the mean ± SD. Statistical analysis using Brown-Forsythe and Welch ANOVA tests followed by Dunnett’s T3 multiple comparisons test. **B**.** *p* = 0.0091 between AAV2/9 empty capsid and AAV2/9-CAG-mNGF groups, * *p* = 0.0397 between AAV2/5 empty capsid and AAV2/5-CAG-mNGF groups, * *p* = 0.0235 between AAV2/9-CAG-mNGF and AAV2/5-CAG-mNGF groups. **C**. ** *p* = 0.0027 between AAV2/9 empty capsid and AAV2/9-CAG-mNGF groups, * *p* = 0.0363 between AAV2/5 empty capsid and AAV2/5-CAG-mNGF groups, * *p* = 0.0127 between AAV2/9-CAG-mNGF and AAV2/5-CAG-mNGF groups. **D**. * *p* = 0.0162 between AAV2/9 empty capsid and AAV2/9-CAG-mNGF groups * *p* = 0.0388 between AAV2/5 empty capsid and AAV2/5-CAG-mNGF groups, * *p* = 0.0368 between AAV2/9-CAG-mNGF and AAV2/5-CAG-mNGF groups. ns, not significant; A. U., arbitrary units. Source data are provided as a Source Data file.

### AAV2/9-mediated gene transfer induces a dose-dependent and long-term protein secretion

Consequently to our results indicating a better protein secretion using AAV2/9 compared to AAV2/5, we chose to focus on the serotype 2/9. To establish the optimal amount of vector genomes (vg) to be injected in LG, we tested three doses and checked the amount of secreted mNGF one month after injection (Figure 6A). While injection of 10^9^ vg did not lead to a drastic increase of the mNGF secretion, a single injection of 10^11^ vg induced the secretion of 75 ng of mNGF per mL of tear film. This high concentration represents a 10-time increase in comparison to physiological mNGF secretion during cornea wound healing process, where a concentration of 7.5 ng / mL was found seven days after abrasion (Figure S3). We concluded that 10^11^ vg / LG was suitable to have a significantly high increase of the protein secretion in the tear film.

**Figure 6.**
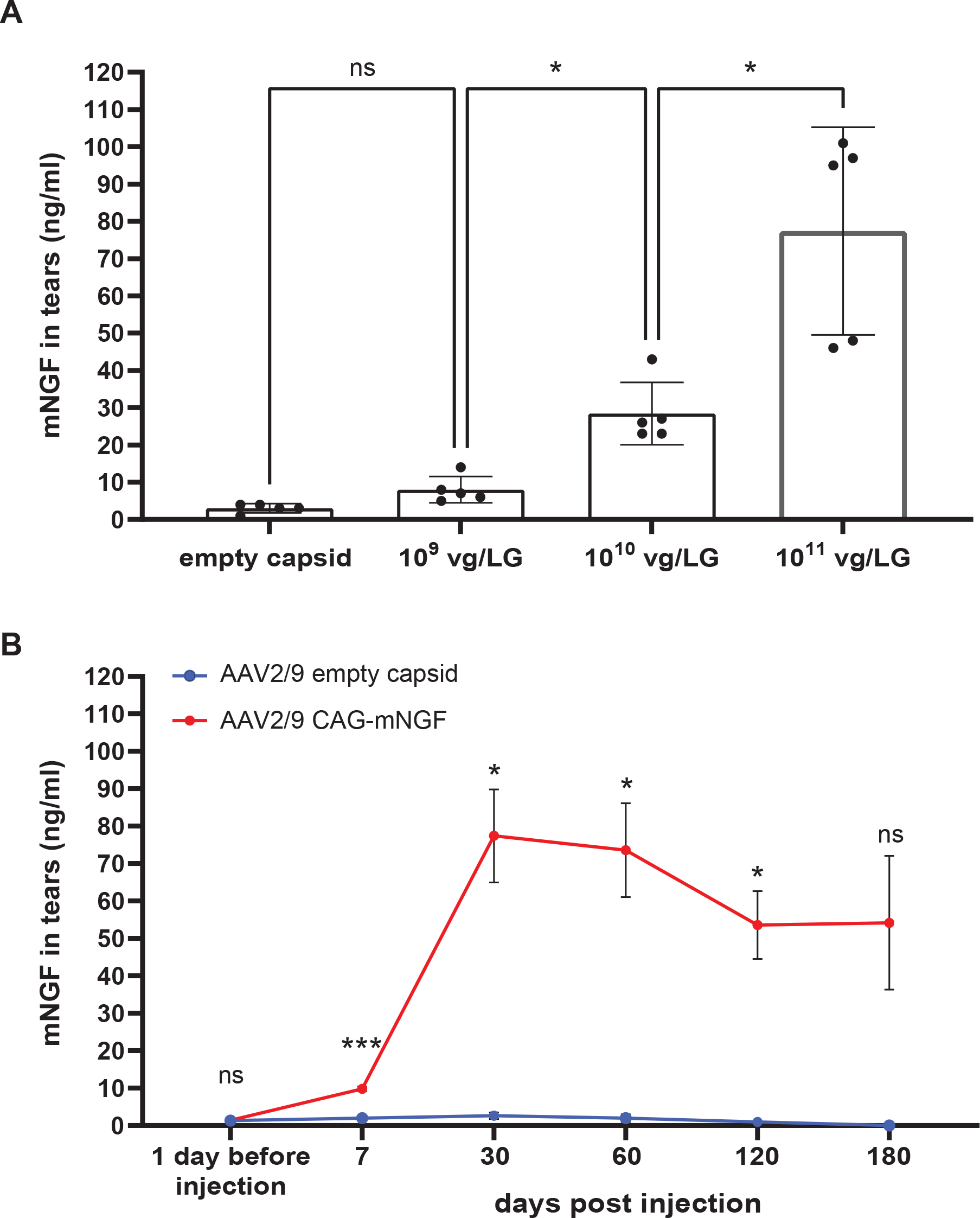
Dose response and kinetic studies after injection of AAV2/9-CAG-mNGF into murine LG reveal a high and long-lasting secretion of mNGF in tears. For the dose response study, mNGF levels were analyzed by ELISA in the tears of AAV2/9-CAG-mNGF-injected mice by increasing dose of vectors (10^9^, 10^10^, and 10^11^ vg / LG in 3 µl, n = 5 mice per dose), when compared to the tears of AAV2/9 empty capsid-injected mice one month post injection. For the kinetic study, mNGF levels were analyzed by ELISA in the tears of AAV2/9-CAG-mNGF-injected mice (10^11^ vg / LG in 3 µl), when compared to the tears of AAV2/9 empty capsid-injected mice, 1 day before injection and at 7, 30, 60, 120 and 180 days post injection (*n* = 3 and 5 for AAV2/9 empty capsid-injected mice and AAV2/9-CAG-mNGF-injected mice respectively). A. Quantification by ELISA of mNGF level (ng / ml) in the tears of injected mice at the indicated doses. Results are expressed as the mean ± SD. Statistical analysis using Brown-Forsythe and Welch ANOVA tests followed by Dunnett’s T3 multiple comparisons test. * *p* = 0.0107 and * *p* = 0.0341 between the doses 10^9^ and 10^10^ vg / LG and between the doses 10^10^ and 10^11^ vg / LG respectively. ns, not significant. B. Quantification by ELISA of mNGF level in tears (ng / ml) at the indicated days post injection. Statistical analysis using repeated measures two-way ANOVA test followed by Sidak’s multiple comparisons test. *** *p* = 0.0009, * *p* =0.0225, * *p* =0.0268, * *p* =0.0260, between AAV2/9 empty capsid and AAV2/9-CAG-mNGF groups at 7, 30, 60, 120 and 180 days post injection respectively. ns, not significant. Source data are provided as a Source Data file.

Subsequently, we evaluated the dynamics of the mNGF secretion over a 6-month period (Figure 6B). The injection of 10^11^ vg into the LG induced a significant increase of the mNGF level found in the tear film already after a week, before reaching a peak at 30 days post injection. The mNGF secretion then decreased to reach a plateau, 120 days after injection with stable expression during the 60 following days. However, although injection of 10^10^ vg/LG of AAV2/9-CAG-mNGF or AAV2/5-CAG-mNG induced a peak 30 days after injection, only the serotype 2/9 induced a significant increase of the mNGF level in the tear film (Figure S4). Nonetheless, the secreted mNGF level for both serotypes decreased and were no more different than the basal level. Taken together, these results show the long-lasting protein secretion when injecting 10^11^ vg/LG, compared to 10^9^ or 10^10^ vg, and confirm that AAV2/9 is more efficient than AAV2/5.

### AAV2/9 vector genome copies in the murine LG correlates with the amount of protein secreted in the tear film

To further analyze further the use of 10^11^ vg / LG, we investigated the biodistribution of the AAV2/9-CAG-mNGF vector after injection into LG. We measured the amount of vector genomes (vg) per diploid genome (dg) in LG, liver and heart, one month after vector injection (Figure 7A). All injected mice showed vector genome copies in LG, indicating that the injection technique is reliable and reproducible. Interestingly, we found around 0.25 vg / dg in the LG, meaning that an average of 1 out of 4 cells was transduced with 1 copy of the AAV2/9-CAG-mNGF vector, which reflects an efficient gene transfer. Despite the detection of the AAV2/9-CAG-mNGF vector genome in liver and heart of injected mice, the levels were very low to negligeable, with 14 and 80-times less vg / dg in liver and heart respectively than in LG.

**Figure 7.**
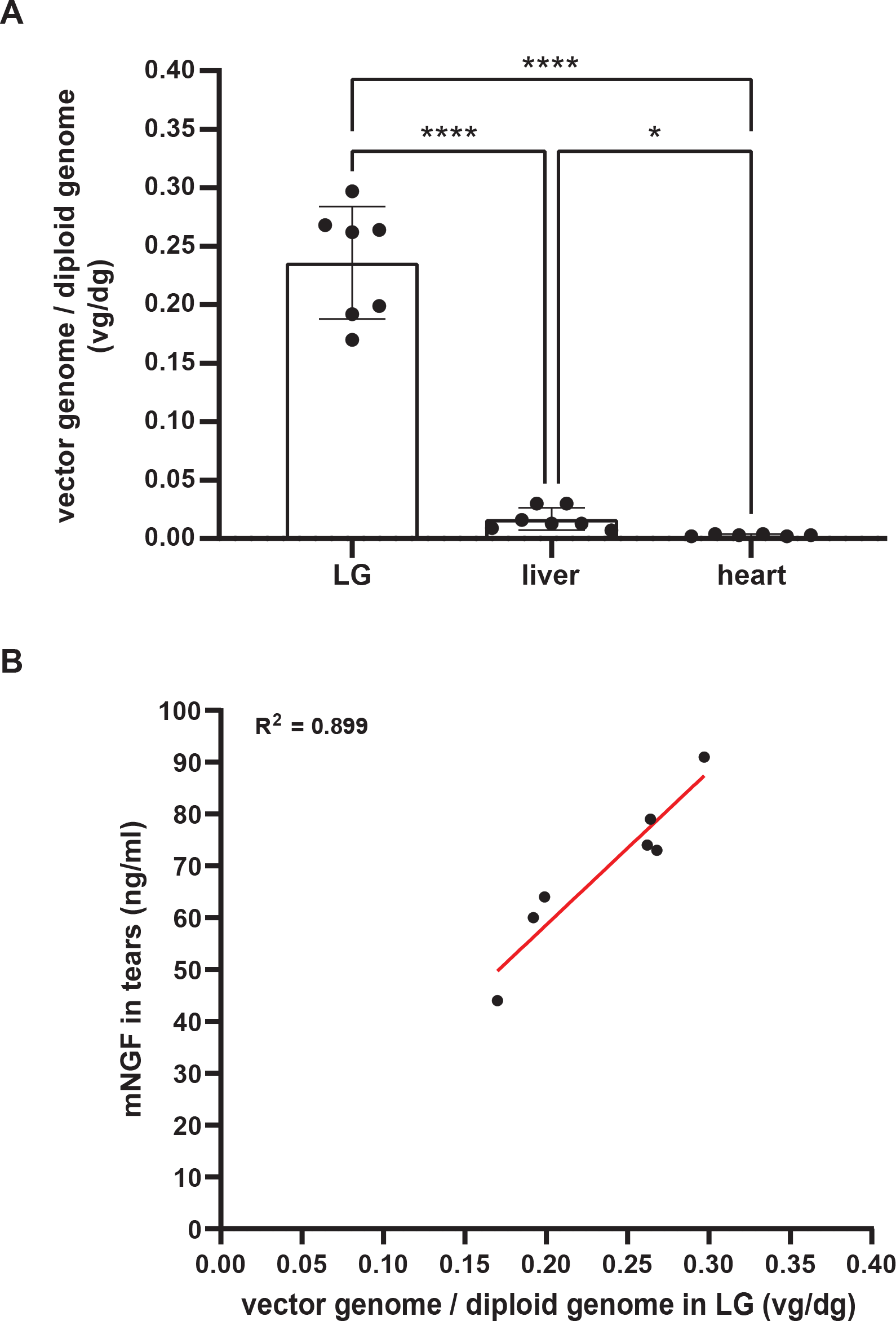
Biodistribution analysis after injection of AAV2/9-CAG-mNGF into murine LG reveals a limited diffusion to peripheral organs. AAV2/9-CAG-mNGF was injected into the murine LG (10^11^ vg / LG in 3 µl, *n* = 7 mice). One month post injection tears were collected and mNGF level was quantified by ELISA. At the same time, mice were sacrificed, LG, liver and heart were collected and analyzed by qPCR. **A.** Quantification of the transduction rate expressed in vector genome / diploid genome (vg / dg) above the lowest limit of quantification (LLOQ) (*n* = 7 for LG and liver samples, *n* = 6 for heart samples). Results are expressed as the mean ± SD. Statistical analysis using Brown-Forsythe and Welch ANOVA tests followed by Dunnett’s T3 multiple comparisons test. **** *p* < 0.0001 between LG and liver, **** *p* < 0.0001 between LG and heart, * *p* = 0.0226 between liver and heart. **B**. Graph showing the statistical linear regression (red line) for each mouse between mNGF level (ng/ml) in tears and vg / dg in LG. R^2^, R squared. Source data are provided as a Source Data file.

Then, we compared the vg / dg values to the concentration of mNGF in the tear film of the same injected mice (Figure 7B). Remarkably, we found a nice correlation between the vg / dg values detected in the LG and the amounts of secreted mNGF in the tears. The increase of vg / dg value corresponded to a higher mNGF secretion.

Taken together, these results confirm the high transduction efficiency displayed by AAV2/9, which is additionally correlated with the amount of protein secreted in the tear film. Moreover, they showed a very limited distribution of AAV2/9 vector to peripheral organs after a single injection into murine LG.

### AAV2/9-mediated mNGF secretion does not affect LG physiology nor corneal integrity

After resolving the parameters for an efficient AAV2/9-mediated mNGF gene transfer, its biodistribution, and induced mNGF secretion, we investigated the impact of the mNGF over secretion on LG physiology and corneal integrity. We injected 10^11^ vg/LG of AAV2/9-CAG-mNGF and measured the impact on tear volume and protein concentration in tears over a period of 6 months. (Figure 8A and 8B). Simultaneously to the increase of mNGF secretion (Figures 4-6), we detected a significant increase of the tear volume from 7 to 120 days post-injection, when compared to the injection of AAV2/9 empty capsid (Figure 8A). Interestingly, this increase of the tear volume was matched with a stability of the protein concentration in tears (Figure 8B). Moreover, to rule out any impact of the injection procedure on LG physiology, we analyzed the tears before and after the AAV2/9 empty capsid injection (Figure S5). Importantly, after injection of the AAV2/9 empty capsid and during the whole 6-month-period, no modification of tear volume nor of total protein concentration was observed (Figure S5). This rule out any impact of the injection procedure itself on LG physiology.

**Figure 8.**
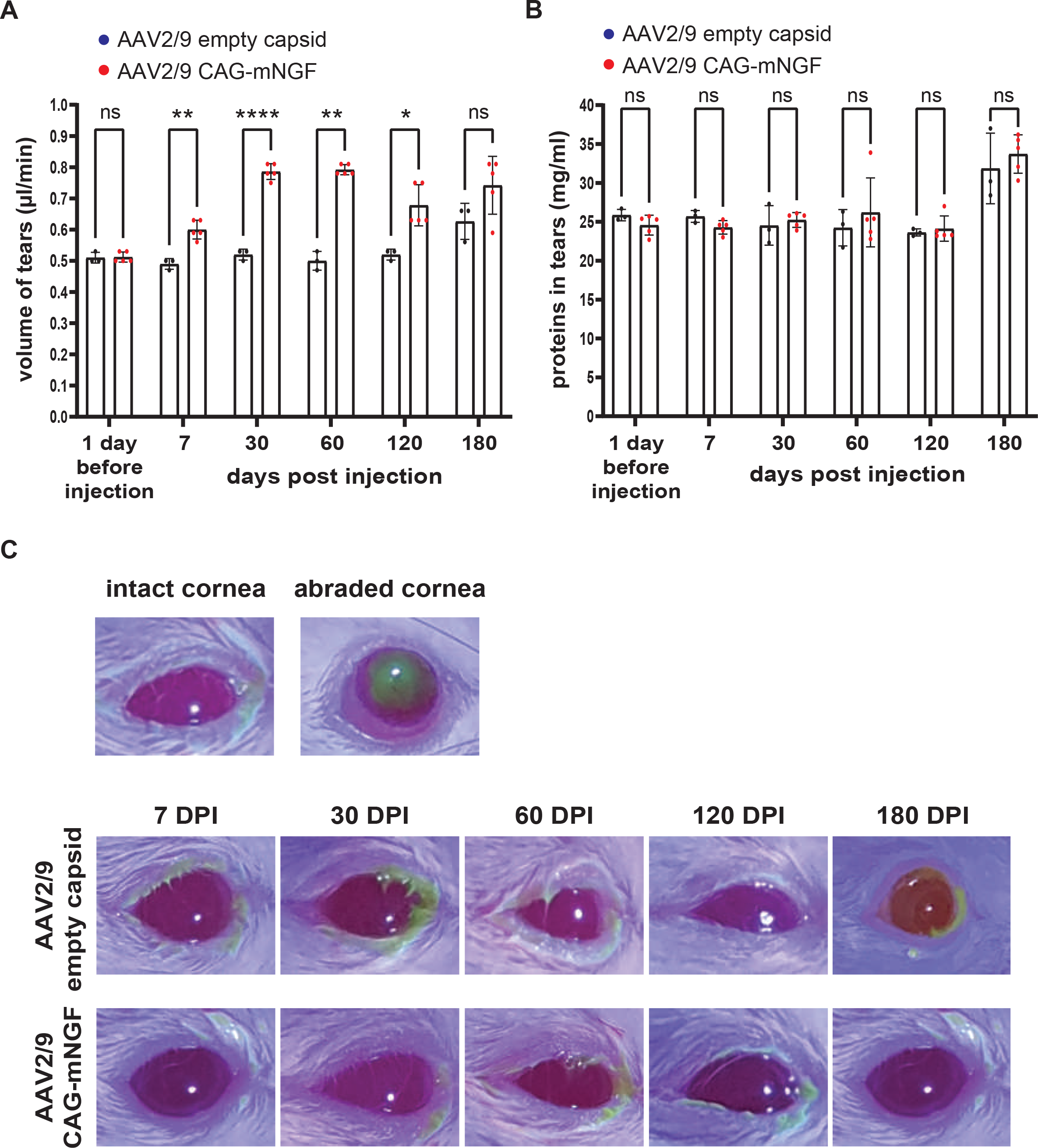
Injection of AAV2/9-CAG-mNGF into murine LG does not alter tear protein concentration nor corneal integrity. AAV2/9 empty capsid or AAV2/9-CAG-mNGF were injected into murine LG (10^11^ vg / LG in 3 µl, *n* = 3 and 5 mice respectively). The volume of tears (**A**), the protein concentration in tears (**B**) and the corneal integrity (**C**) were assessed 1 day before injection and 7, 30, 60, 120 days and 180 post injection. **A.** Graph showing the volume of tears (µl / min) collected before injection and at the indicated days post injection. Results are expressed as the mean ± SD. Statistical analysis using repeated measures two-way ANOVA test followed by Sidak’s multiple comparisons test. ** *p* = 0.0036, **** *p* < 0.0001, ** *p* = 0.0057, * *p* = 0.0246 between AAV2/9 empty capsid and AAV2/9 CAG-mNGF groups at 7, 30, 60 and 120 days post injection respectively. ns, not significant. **B.** Graph showing the protein concentration in tears (mg / ml) collected before injection and at the indicated days post injection. Results are expressed as the mean ± SD. Statistical analysis using repeated measures two-way ANOVA test followed by Sidak’s multiple comparisons test. ns, not significant. **C**. Eye images obtained from the fluorescein stain test. Fluorescein signal highlights an abraded cornea in bright green, while an intact cornea remains dark. Source data are provided as a Source Data file.

In addition, we used fluorescein staining to monitor the corneal epithelium integrity and visualize any adverse effect that the injection procedure and the tear film modification could have on the cornea. Fluorescein stains areas where the corneal epithelial barrier is defected^1^, as after corneal abrasion (Figure 8C). During the 180 days after AAV2/9 injection, we never observed any fluorescein staining, whether we injected an empty capsid, or induced mNGF over secretion. We concluded that AAV2/9 injection does not impact the corneal epithelium integrity.

To investigate further the effect of the mNGF over secretion on cornea, we visualized corneal innervation with βIII tubulin immunolabelling, which is a pan neuronal marker. We showed that the β gross morphology of corneal fibers was not affected by the over secretion of mNGF in the tear film (Figure 9A). Furthermore, we performed von Frey tests on the corneas of injected mice. We demonstrated that the constant over secretion of mNGF did not modify corneal sensitivity (Figure 9B). All together, these results demonstrate that the injection procedure and mNGF over secretion respect the LG physiology and the corneal integrity.

**Figure 9.**
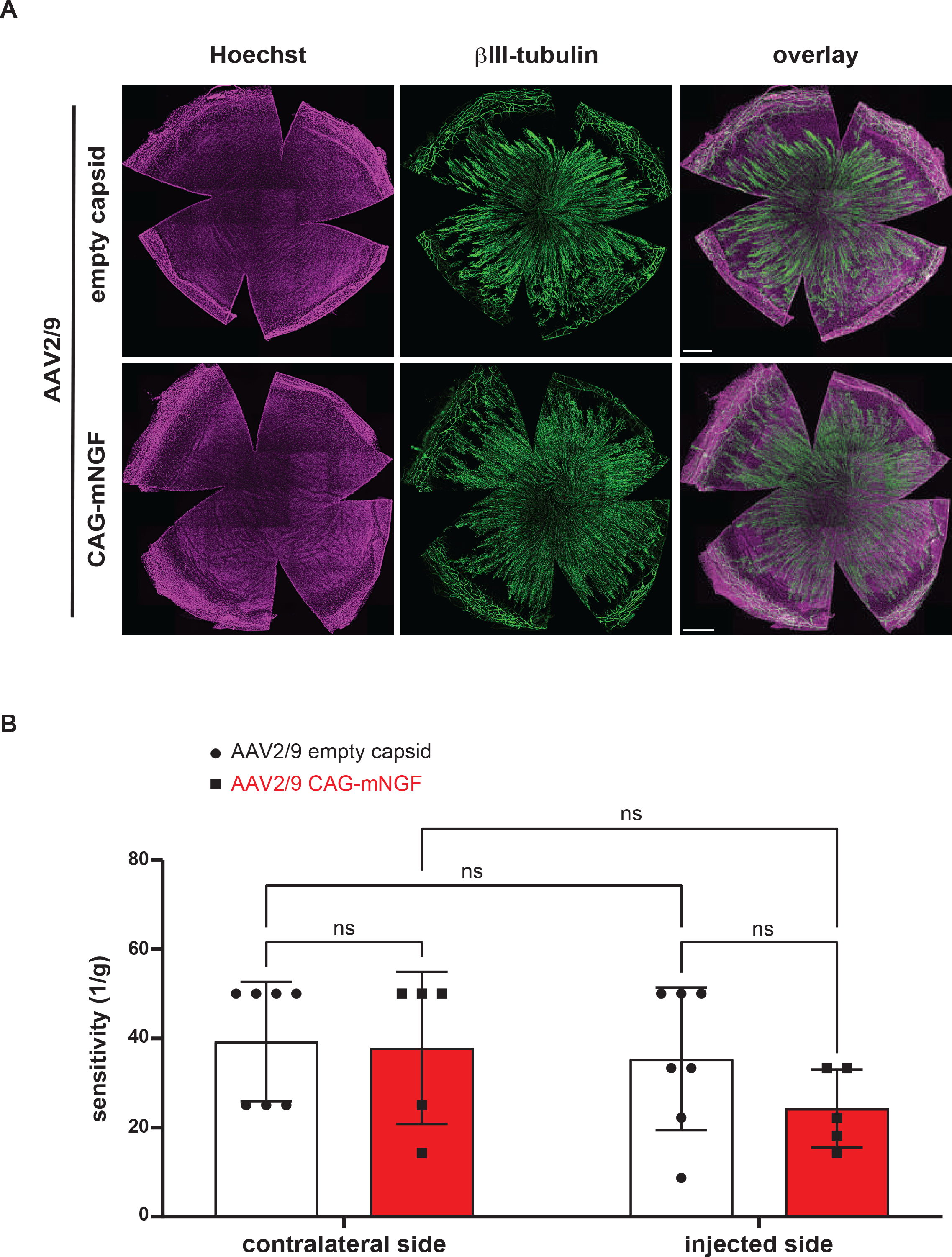
Injection of AAV2/9-CAG-mNGF into murine LG does not affect corneal innervation nor sensitivity. **A**. Representative images showing corneal innervation. Whole corneas were immunostained for III-tubulin (green) following a single injection of AAV2/9-CAG-mNGF into β murine LG, when compared to AAV2/9 empty capsid-injected mice (10^11^ vg / LG in 3 µl, *n* = 3 mice per group). Mice were sacrificed two weeks post injection. Nuclei are counterstained with BioTracker NIR694 (magenta). Scale bar: 500 µm. **B**. Quantification of the corneal sensitivity (1 / g) measured by von Frey test performed both on the contralateral side and the injected side of AAV2/9 empty capsid (white plots, *n* = 7) or AAV2/9-CAG-mNGF (red plots, *n* = 5)-injected mice. Von Frey analysis was performed two weeks post injection. Results are expressed as the mean ± SD. Statistical analysis using two-way ANOVA test followed by Sidak’s multiple comparisons test. ns, not significant. Source data are provided as a Source Data file.

## Discussion

Among the sight threatening diseases, corneal defects are the fourth cause for blindness globally. Except for physical harm, corneal blindness often results from a combination of intrinsic and extrinsic causes. In the case of genetic diseases, such as aniridia^29, 30^, or systemic diseases, such as diabetes^13^, corneal defect is often associated with corneal microenvironment dysregulation^31^.

The aim of this study was to establish an innovative and attractive strategy to tackle corneal defects by sustainably modulating the tear film composition. For this purpose, we chose an AAV-mediated gene transfer to use LG as a bioreactor producing specific transgenes. Nevertheless, several major variables are crucial to achieve an efficient transduction of the target cells, and thus require to be well-designed^17, 32^. First, the route of administration determines both the efficacy and the biosafety pattern of an AAV-based gene transfer. In this study, we used a local injection into LG as this organ is easily accessible by surgery. Moreover, we showed that a local injection allows concentration and time residence of AAV vectors in vicinity to the target cells, limits the biodistribution of the vector to non-targeted tissues and thus the risk of toxicity. For instance, local injection is well-established for ocular diseases, particularly in retinal disorders, as several anatomical sites are commonly targeted: subretinal, intravitreal, intracameral, suprachoroidal and topical^33, 34^. Furthermore, the injection method is a key parameter to reach a large diffusion in the target organ. While a systemic injection leads to large diffusion in all the organism including the target organ^35^, when using local injection, the development of specific injection procedures are necessary. For example, several injections at precise coordinates are required for a large diffusion in the brain^36^. Interestingly, a pneumatic picopump system applying multiple short-time pressure pulses has been reported to transduce the whole sciatic nerve of rodents^37^. In this study, we used 34-gauge beveled needle linked to a 10- μL Hamilton syringe to perform injections into LG. We injected 3 µl of AAV vectors by following the path of the LG main duct. This procedure of injection led to a large diffusion in the gland, and showed an efficient gene transfer in all epithelial cell types in the murine LG.

The promoter used to drive the expression of the transgene is a second parameter determining the transduction efficiency and the biosafety pattern of the injection^17, 32^. Indeed, the promoter controls the transgene expression level. Commonly, AAV-mediated gene transfer uses an ubiquitous and strong promoter, such as CAG and CMV, to achieve high transgene expression^38, 39^. Moreover, researchers usually employ such promoters when trying to establish a proof of concept for AAV-mediated gene transfer into a specific organ. In this study, we utilized a CAG promoter to provide a proof of concept of the tear fluid composition modulation after an AAV-mediated gene transfer into murine LG. On the other hand, a high transgene expression level is not always desired. For instance, transgene over expression above physiological levels has been reported to be toxic^40^. This toxicity must be correlated to the serotype of the AAV vector. Indeed, the serotype represents another parameter influencing both the transduction efficiency and the biosafety of an injection as each AAV serotypes exhibit different cell and tissue tropisms^41^. Therefore, the combination of the route of administration, the AAV serotype, the promoter driving the transgene expression and the dose of vector have to be carefully designed as it may induce transgene expression in off-target tissues and thus lead to dramatic toxicities^42^. In this study, we compared the AAV serotypes 2/5 and 2/9 to first transduce the murine LG efficiently and then to modulate the tear film composition. Notably, while both serotypes led to an efficient gene transfer in the LG, AAV2/9 gave a better yield of GFP production and mNGF secretion compared to AAV2/5. Interestingly, although the difference of transduction efficiency was reported previously^24^, the 2 to 4 time fold in mNGF secretion is startling. Indeed, until now, no study has reported a transgene secretion in the tear fluid after an AAV-mediated gene transfer into LG. Furthermore, we tested increasing dose of AAV2/9 to evaluate the mNGF secretion in the tear film. Interestingly, while 10^11^ vg / LG dose led to the highest mNGF secretion, 10^10^ vg / LG was sufficient to induce a significant secretion. Given that we used the serotype 2/9, known to show high heart and liver tropism in rodent^43^, and a strong CAG promoter, we performed a biodistribution study for AAV2/9. This study was paramount to determine the potential off-targets of our strategy, which may lead to both unwanted toxicity and immunogenicity. At the highest injected dose of 10^11^ vg / LG, we showed an average of 0.25 vg / dg in LG, only 0.017 vg / dg in liver and 0.003 vg /dg in heart of injected mice, which corresponded to 14- and 80-times less vg / dg than in LG respectively. Importantly, these levels of AAV2/9 found in liver and heart were very low to negligible, from 100 to 10 000 times lower than those obtained after intravenous^39^ or intrathecal^44^ injections.

Different strategies exist to limit the transgene expression in off-targets and its related adverse effects, including the use of cell-type specific promoters^39^ or the incorporation of miRNA binding sites in the AAV gene expression cassette^45^. We can speculate that the use of a 10-time lower dose of AAV2/9 vector, providing a high secretion of mNGF, could bring a lower off-target transduction. Injection of 10^10^ vg / LG of AAV led to a faster secretion of mNGF with AAV2/9 compared to AAV2/5. After a peak 30-day post-injection, the mNGF secretion decreased to the physiological level with both serotypes. However, injection of 10^11^ vg / LG results in over secretion of mNGF during several months. This is an important aspect for the subsequent use of this method. While a lower dose could be used for transitory pathologies, such as corneal graft, or recurrent corneal abscesses, a higher dose could be of interest for chronical pathologies, such as neurotrophic keratitis, or dry eye diseases.

The long-term secretion of mNGF is a remarkable discovery, which must be linked to the LG physiology. As all epithelial organs, such as skin, cornea or mammary glands, the epithelial compartment contains stem cells regenerating continuously the organ, and healing it if necessary^46^. The AAV vector is a non-integrative vector that can be lost when cell division occurs^47^. Therefore, only non-proliferating cells will keep the transgene and lead to protein secretion. Our results demonstrate that gene therapy is successful on this epithelial organ and highlighting the low turn-over of epithelial cells in the LG. Knowing that 30 days after injection, there was around one AAV2/9 vector genome in every 4 cells, it would be of great interest to monitor over a long period of time to see which cells retain the AAV2/9 genome. This method would give a deeper understanding of LG fundamental biology. Of course, this long term follow-up could only be done at the cellular level, as with age, the LG physiology is randomly affected in individuals^8^. Therefore, measuring tear volume and tear protein content could be misleading because of aging impact more than AAV2/9 injection. Nevertheless, we demonstrated that up to 6 months after injection into the murine LG, our approach has no detrimental impact on the LG physiology.

The presence of mature mNGF in the tear film is important for its functionality. We showed that in the absence of mNGF over secretion, only a negligible part of the pro-mNGF was processed into its mature form. Interestingly, the mNGF over secretion led to the presence of mature mNGF in the tears. However, we cannot explain why after AAV2/9 injection, 57 % of the total mNGF is matured, and only 34 % after AAV2/5 injection. We can only hypothesize a threshold effect leading to a better pro-mNGF processing into mature mNGF when there is an increase of secretion in the tear film.

Despite the large amount of mNGF and its constant presence on the cornea, we did not detect any impact on corneal innervation, nor on corneal sensitivity. We hypothesize that the robustness of corneal innervation maintains the system under control in physiological conditions, and mNGF alone is not sufficient to disturb this system. Most likely, under pathological conditions, i.e., physical harm, or neurotrophic keratopathy, the system would be sufficiently perturbed to visualize the effect of mNGF on corneal innervation.

Collectively, our results show that LG gene therapy could be established to modify specifically the tear film to support corneal physiology. The main challenge in using AAV vectors for epithelial organ gene therapy is the high renewal rate of epithelial cells. Here, we demonstrate that there is a non-renewing cell population in the LG that retains the secretory capabilities to produce a large amount of mNGF for over 6 months. The long term over secretion of mNGF could replace the use of NGF supplemented eyedrops, used to treat neurotrophic keratitis, such as observed in diabetes^12^ or neurodegenerative diseases^48^. Notably, by substituting mNGF with another gene, other corneal defects could be treated.

## Material and Methods

### Study design

The goal of this study was to assess the transduction pattern of AAV vector serotypes 2/5 and 2/9 after a single injection into murine LG. Then, to evaluate the efficiency and the safety of a AAV2/9- mediated gene transfer of mNGF in the murine LG, for its secretion in the tear film. The main readouts of this study included: the transduction pattern analyzed by IHC and Western blot, the secretion of mNGF measured in tears by ELISA and Western blot, the AAV2/9 biodistribution by qPCR, the injection biosafety by analyzing the corneal integrity and innervation, the volume of tears and the protein concentration in tears. Experimental groups were sized according to the literature to allow statistical analysis. No outliers were excluded from the study, except mice exhibiting spontaneous eye damages after the surgery or during the experiments. Behavioral data obtained from animals displaying eye damages unrelated to the abrasion procedure during the study were excluded. Scientists who performed the experiments and analysis were blinded to the group’s identity. Data were analyzed by those carrying out the experiments and verified by the supervisor.

### Cloning and vector production

The recombinant AAV vectors developed were called AAV hereafter. Cloning of the enhanced GFP (GFP) and the mNGF in pAAV and AAV vector productions were provided by the vector core of the TarGeT (Translational Gene Therapy) Laboratory of Nantes, INSERM UMR 1089 (Nantes University, France). Briefly, single-stranded AAV2/5 and AAV2/9 CAG-GFP, single-stranded AAV2/5 and AAV2/9 CAG-mNGF vectors were obtained from pAAV CAG-GFP and pAAV CAG- mNGF plasmids respectively, containing AAV2 inverted terminal sequences, CAG promoter and BGH polyA signal. AAV2/5 and AAV2/9 CAG-GFP vectors were used to assess the transduction pattern after injection in the murine LG. AAV2/5 and AAV2/9 CAG-mNGF vectors were used to evaluate the efficiency of an AAV-based gene transfer in the LG allowing mNGF secretion in the tear fluid. Single-stranded vectors containing AAV2/5 and AAV2/9 empty capsid served as controls.

Vector production was performed following the protocol of vector core of the TarGeT Laboratory of Nates^49^. Briefly, recombinant AAVs were manufactured by co-transfection of HEK293 cells and purified by cesium chloride density gradients followed by extensive dialysis against phosphate-buffered saline (PBS). Vector titers were determined by qPCR and expressed as vector genome (vg) / ml. The target amplicons correspond to the inverted terminal repeat (ITR) sequences, ITR-2 (ITR-2 Forward: GGAACCCCTAGTGATGGAGTT, ITR-2 Reverse: CGGCCTCAGTGAGCGA, Taqman Probe used for vector titer: FAM- CACTCCCTCTCTGCGCGCTCG-BBQ).

### Animals included in this study

All mice experiments were approved by the local ethical committee and the “ministère de la recherche et de l’enseignement supérieur” (authorization 2016080510211993 version2). All the procedures were performed in accordance with the French regulation for the animal procedure (French decree 2013-118) and with specific European Union guidelines for the protection of animal welfare (Directive 2010/63/EU). Mice were maintained on a 12 h dark, 12 h light cycle with a humidity between 40 and 60 % and an ambient temperature of 21-22 °C.

### Surgery and Vector delivery

Twelve-week-old Swiss/CD1 female mice (Janvier Labs, France) were injected into the right LG. The LG injection of AAV vectors was performed under anesthesia with a mixture of Ketamine (70 mg / kg, Imalgene® 1000, Centravet, France) and Medetomidine (1 mg / kg, Domitor®, Centravet, France). One drop of Ocry-gel (Centravet, France) was applied to each eye. The skin on the cheek under the right ear was disinfected with vetedine solution (Centravet, France) and ethanol 70% and then cut above the LG location. Next, the viral solution was injected using a 34-gauge beveled needle (Hamilton, reference 207434, Reno, NV, USA) linked to a 10-μl Hamilton syringe (1701 RN serie, Hamilton, reference 7653-01, Reno, NV, USA). Wound were closed by suture wires (Novosyn® 6/0, reference C0068006, B Braun) and then disinfected with vetedine solution (Centravet, France). After surgery, mice were treated with Buprenorphine (100 μ Brupecare®, Centravet, France) and were woken up with Atipamezole (1 mg / kg, Antisedan®, Centravet, France).

AAV vector solutions were prepared by diluting vectors at the right titer with sterile PBS and 0.01 % of Fast Green (Sigma-Aldrich, reference F7252, France). For all experimental studies, mice were unilaterally injected in the right LG with 3 µl of vectors indicated below. For the transduction pattern study, AAV2/5 and AAV2/9 CAG GFP were injected at 10^10^ vg / LG. For the dose response study, AAV2/9 CAG-mNGF was injected at 10^9^, 10^10^ or 10^11^ vg / LG. For the kinetic study, AAV2/5 CAG-mNGF was injected at 10^10^ vg / LG and AAV2/9 CAG-mNGF was injected at 10^10^ and 10^11^ vg / LG. For the biodistribution and the biosafety studies, AAV2/9 CAG-mNGF was injected at 10^11^ vg / LG. Animals injected with AAV2/5 or AAV2/9 empty capsid served as control.

### Corneal abrasion

Corneal abrasions were performed as previously described^1,3, 50^. Briefly, an ocular burr (Algerbrush II, reference BR2-5 0,5 mm, Alger company) was used on mice that were anesthetized with a mixture of Ketamine (70 mg / kg, Imalgene® 1000, Centravet, France) and Medetomidine (1 mg / kg, Domitor®, Centravet, France). Abrasions were performed unilaterally. A fluorescein solution (1% in PBS, Sigma-Aldrich) was used to visualize the wound under a cobalt blue light. After abrasion, one drop of Ocry-gel (Centravet, France) was applied to each eye, mice were treated with Buprenorphine (100 μg / kg, Brupecare®, Centravet, France) and were woken up with Atipamezole (1 mg / kg, Antisedan®, Centravet, France).

### Tissue collection and processing

Tears were collected using a 1 µl-microcapillary (Sigma-Aldrich, reference P1424, France) for one minute, 1 day before injection and 30 days post injection for the dose response and biodistribution studies; 1 day before injection and 7, 30, 60, 120 and 180 days post injection for the kinetic and biosafety studies; 1 day before abrasion and 1, 3 and 7 days post abrasion for the corneal abrasion study.

For the transduction pattern and the biodistribution studies, mice were euthanized 30 days post injection using pentobarbital (54.7 mg/mL, 140 mg/kg, Centravet, France). They were transcardially perfused with sterile PBS and tissues were quickly dissected. Tissues were then fixed for 45 min in 4% paraformaldehyde solution (AntigenFix, Diapath, reference P0014, France) at room temperature or directly snap-frozen in liquid nitrogen and stored at -80°C for IHC and molecular/biochemical analysis respectively.

For whole cornea imaging, mice were euthanized by cervical dislocation, enucleated with curved scissors by cutting the optic nerve. Collected eyes were then fixed for 20 min in 4% paraformaldehyde solution (AntigenFix, Diapath, reference P0014, France) at room temperature. After PBS washes, eyes were dehydrated during 2 h in 50% ethanol/PBS and then stored at 4°C in 70% ethanol/PBS.

### Tear volume, protein concentration and mNGF analyses in tears

The volume of tears per minute (µl / min) and the protein concentration in tears (mg / ml) using the BCA protein assay kit (Fisher Scientific, reference 10678484, France) were measured and expressed as the mean ± SD.

The mNGF level in tears upon a AAV2/9 CAG-mNGF injection in the LG was dosed using the mNGF ELISA kit (Sigma-Aldrich, reference RAB1119, France) following the manufacturer’s recommendations. Measures were performed on a CLARIOstar microplate reader (BMG Labtech, France), and analyzed using the CLARIOstar software (version 5.60 R2). Results are expressed as the mean ± SD.

### IHC study

The following antibodies were used for IHC studies: chicken anti-GFP (Aves Labs, reference GFP- 1020, 1/1000), mouse anti-Ecadh (BD Biosciences, reference 610182, 1/300), rabbit anti-Krt19 (Abcam, reference ab52625, 1/200), rabbit anti-βIII tubulin (Abcam, reference ab18207, 1/1000), goat anti-chicken Alexa Fluor 488 (Thermo Fisher Scientific, reference A-32931, 1/500), goat anti-mouse Alexa Fluor 568 (Thermo Fisher Scientific, reference A-11004, 1/500) and goat anti-rabbit

Alexa Fluor 568 (Abcam, reference ab175471, 1/500), goat anti-rabbit Alexa Fluor 488 (Abcam, reference ab11008, 1/500) and goat anti-rabbit Alexa Fluor 568 (Abcam, reference ab175471, 1/500). Nuclei were counterstained with Hoechst 33342 (Thermo Fisher Scientific, reference H3570, 1/2000) and BioTracker NIR694 (Merck, reference SCT118, 1/400).

### Immunohistochemistry on LG frozen sections

Following fixation, LG were incubated 24 h in two successive baths of 6 % and 30 % sucrose and then embedded in Optimal Cutting Temperature (OCT tissue freezing medium, MM France, reference F/TFM-C) and stored at -80 °C. Longitudinal sections (10 µm of thickness) were cut using a cryostat apparatus (LEICA CM3050). For primary antibodies produced in rabbit and chicken, cryosections were blocked with a mixture of 5 % Goat Serum (GS, Thermo Fisher Scientific, reference 16210064), 5 % of fish skin gelatin (FSG, Sigma-Aldrich, reference G7765) and 0.1 % triton X-100 in PBS for 1 h at room temperature. For primary antibodies produced in mouse (mouse anti-Ecadh), the blocking step described above was followed by an incubation with goat anti-mouse immunoglobulins (Abcam, reference ab6668, 1/200) during 1 h at room temperature. Cryosections were then incubated overnight at 4 °C with primary antibodies diluted in GS/FSG/Triton/PBS mixture, washed three times with 0.1 % triton X-100 / PBS and subsequently incubated 1 h at room temperature with secondary antibodies diluted in GS / FSG / Triton / PBS mixture. After several PBS washes, cryosections were mounted in Fluoromount-G mounting medium (Invitrogen, reference 00-4958-02).

## Immunohistochemistry on whole cornea

Eyes were rehydrated in 50 % ethanol/PBS during 2 h and washed twice in PBS for 15 minutes at room temperature. Corneas were dissected and permeabilized with 0.5% Triton-X-100/PBS on a rocking agitator for 1 h and then blocked in 5 % GS (Thermo Fisher Scientific, reference 16210064) 2.5 % FSG (Sigma-Aldrich, reference G7765) in 0.1 % Triton X-100/PBS at room temperature. Corneas were incubated in primary antibody diluted in blocking solution overnight at 4°C on a rocking agitator and rinsed in 0.1 % Triton X-100/PBS at room temperature (three times 1 h). Next, samples were incubated with secondary antibodies as previously mentioned. After the washes, nuclei were stained 10 minutes with BioTracker NIR694 (Merck, reference SCT118) and washed in PBS. Corneas were cut at four corners and mounted in Fluoromount-G mounting medium (Invitrogen, reference 00-4958-02), epithelium facing the coverslip.

### Imaging

LG images were acquired using Zen Black software (version 2.3 SP1, Zeiss, France) on a LSM 880 confocal microscope (Zeiss, France). Whole LG section images were obtained using a 20x / 0.8 objective while co-immunostaining of GFP with E-cadherin or Keratin19 proteins were observed via 0.36 µm step size z-stacks using a 63x / 1.4 oil immersion objective. Images were then processed with Zen Black software (version 2.3 SP1, Zeiss, France) and Zen Blue lite software (version 3.2, Zeiss, France).

Whole cornea images were acquired using the navigator module on a Leica Thunder Imager Tissue microscope with Large Volume Computational Clearing (LVCC) process. Images were obtained using a 20x/0.55 objective with LAS X software (3.7.4) and processed with Imaris Bitplane software (version 9.8.0).

### Western blot

Frozen LG were crushed with a pestle and mortar pre cooled at -80°C, solubilized in Pierce® RIPA lysis buffer (Thermo Fisher Scientific, reference 89900, France) supplemented with protease inhibitors (Halt™ Protease Inhibitor Cocktail, Thermo Fisher Scientific, reference 87786, France), homogenized on a rotating wheel at 4 °C overnight and then centrifuged at 16 900 g (Centrifuge 5418R, Eppendorf) for 30 min at 4 °C. Supernatants were recovered and protein concentration of LG lysates or collected tears were quantified using the BCA protein assay kit (Thermo Fisher Scientific, reference 10678484, France). Fourty and 8 µg of proteins from LG lysates and collected tears respectively were loaded on any kD precast polyacrylamide gels (Mini-Protean® TGX™ gels, Bio Rad, reference 4568124, France). Proteins were transferred to nitrocellulose membranes (Trans-Blot Turbo Mini 0.2 µm Nitrocellulose Transfer Pack, Bio Rad, reference 1704158, France) through semi-dry transfer process (Bio Rad Trans-Blot Turbo system). Membranes were incubated with the REVERT™ total protein stain solution (LI-COR Biosciences, reference 926-11015, France) to record the overall amount of protein per well. They were then blocked for 1 h at room temperature using Intercept® blocking buffer (LI-COR Biosciences, reference 927-60001, France). They were incubated with the following primary antibodies overnight at 4 °C in LI-COR blocking buffer: chicken anti-GFP (Aves Labs, reference GFP-1020, 1/2000) or rabbit mNGF (Abcam, reference Ab52918, 1/500). Following three washes with TBS containing 0.1% Tween (TBST) for 15 min, secondary antibodies were incubated at a 1/15000 dilution in LI-COR blocking buffer: donkey anti chicken IR Dye 800CW (LI-COR Biosciences, reference 926-32218) or donkey anti rabbit IR Dye 800CW (LI-COR Biosciences, reference 926-32213). After three washes in TBST for 15 min, images were acquired with an Odyssey CLX LI-COR Imaging System (LI-COR Biosciences, France) and the quantifications were performed with Image Studio lite software (version 5.2). The GFP and the mNGF protein levels were both normalized to the total amount of protein loaded per well. The results are expressed as the mean ± SD.

### AAV2/9 biodistribution study

LG, liver and heart from AAV2/9-CAG-mNGF-injected mice (10^11^ vg / LG, *n* = 7) were collected one month post injection in DNA-free, RNAse/DNAse-free and PCR inhibitor-free certified microtubes. Tissue samples were collected immediately after sacrifice, snap-frozen in liquid nitrogen and stored at -80 °C in conditions that minimize cross-contamination and avoid qPCR inhibition. Extraction of genomic DNA (gDNA) from tissues using the Gentra Puregene kit (Qiagen, reference 158445, France) and Tissue Lyser II (Qiagen, reference 85300, France) was performed in accordance with the manufacturer’s recommendations. Vector genome copy number was determined using a primer/FAM-TAMRA probe combination designed to amplify a specific region of the *BGH* transgene (*BGH* forward: TCTAGTTGCCAGCCATCTGTTGT, *BGH* reverse: TGGGAGTGGCACCTTCCA, *BGH* probe: FAM-TCCCCCGTGCCTTCCTTGACC-TAMRA). qPCR analyses were conducted on a StepOne Plus apparatus (Applied Biosystems®, Thermo Fisher Scientific, France) using 50 ng of gDNA in triplicates and the following cycling conditions: denaturation step (20 sec, 95°C) followed by a total of 45 cycles (1 sec, 95°C; 20 sec, 60°C). All reactions were performed in a final volume of 20 µl containing template DNA, Premix Ex Taq (Takara/Ozyme, reference RR390L, France), 0.4µl of ROX reference Dye (Takara/ozyme, reference RR390L, France), 0.2 µmol/l of each primer and 0.1 µmol/l of Taqman® probe (Dual-Labeled Probes,Sigma-Aldrich, France). Endogenous gDNA copy numbers were determined using the following primers/FAM-BHQ1 probe combination, designed to amplify a specific region of the murine *Albumin* sequence (murine *Albumin* forward: AACTGAAACTTTGGGAGGT, murine *Albumin* reverse: GGAGCACTTCATTCTCTGAC, murine *Albumin* probe: FAM- AGCTTGATGGTGTGAAGGAGAAAG-BHQ1). qPCR analyses were conducted on a C1000 touch thermal cycler (Bio-Rad, France) using 50 ng of gDNA in triplicates and the following cycling conditions: denaturation step (20 sec, 95°C) followed by a total of 45 cycles (3 sec, 95°C; 30 sec, 60°C). All reactions were performed in a final volume of 20 µl containing template DNA, Premix Ex Taq (Takara/Ozyme, reference RR390L,France), 0.25 µmol/l of each primer and 0.2 µmol/l of Taqman® probe (Dual-Labeled Probes,Sigma-Aldrich, France). For each sample, threshold cycle (Ct) values were compared with those obtained with different dilutions of linearized standard plasmids (containing either the *BGH* expression cassette or the murine *Albumin* gene) using Bio-Rad CFX Maestro 2.2 software (version version 5.2.008.0222) or StepOne software (version 2.3) for the C1000 touch thermal cycler and the StepOne Plus apparatus respectively. The absence of qPCR inhibition in the presence of gDNA was checked by analyzing 50 ng of gDNA extracted from tissue samples from two AAV2/9 empty capsid-injected control mice. Results were expressed in vector genome copies per diploid genome (vg / dg) as the mean ± SD. The lowest limit of quantification (LLOQ) was determined as 0.001 vg/dg. Only values of vg / dg above the LLOQ were presented.

### Von Frey test

Corneal sensitivity was evaluated using von Frey filaments (Bioseb, reference bio-VF-M) as previously described^51^. Filaments with define forces from 0.008 to 0.6 g were applied on the cornea of an immobilized mouse until an eye-blink reflex was observed. Mice were habituated every day for five days. Von Frey test was performed on each cornea for two consecutive days (contralateral side and injected side). The values obtained from these two days were averaged for each cornea. As the values were represented in g, we displayed them as 1/g to reflect the sensitivity. Results are expressed as the mean ± SD. Behavioural experiments and analysis were performed by the same experimenter in single-blinded conditions throughout the study.

### Statistical analysis

Data were analyzed with GraphPad Prism software (version 9.1.2, Prism, CA, USA) and expressed as the mean ± SD as indicated in the Figure legends. Statistical differences between mean values were tested using Brown-Forsythe and Welch ANOVA tests followed by Dunnett’s T3 multiple comparisons test, repeated measures two-way ANOVA test followed by Sidak’s or Tukey’s multiple comparisons test, repeated measure one-way ANOVA test followed by Dunnett’s comparisons test, Friedman’s one-way ANOVA test followed by Dunn’s multiple comparisons test or simple linear regression as indicated in the Figure legends. Differences between values were considered significant with: *p < 0.05, ** p < 0.01, *** p < 0.001, **** p < 0.0001. All exact p-values are available in the Source Data File.

### Data and materials availability

All data and materials are available upon request. The source data underlying Figures 3-7, 8A, 8B, 9B and Supplementary Figures S3-S5 are provided as a Source Data File.

## Ethics declarations

The authors declare no competing interests.

## Author contributions

Conceptualization: B.G., F.M.; Methodology: B.G., L.M., L.H.; Validation: B.G., F.M.; Formal analysis: B.G., L.H., F.M.; Investigation: B.G., L.M., C.A., L.H., N.F., A.K., A.P., C.L.G., V.B., F.M. Data analysis: B.G., F.M.; Writing-Original Draft: B.G., F.M.; Writing-Review and Editing: all authors; Supervision: F.M.; Project administration: F.M.; Funding acquisition: C.D., F.M.

## Acknowledgments

The authors thank the different technical platforms of the Institute for Neurosciences of Montpellier, especially the RAM-Neuro, animal core facility supervised by Denis Greuet, the imaging facility MRI, member of the national infrastructure France-BioImaging infrastructure supported by the French National Research Agency (ANR-10-INBS-04, «Investments for the future»), and the CPV vector core and Preclinical Analytics Core from the TarGeT lab, INSERM UMR 1089, Nantes University (https://umr1089.univ-nantes.fr/facilities-cores/). This research was supported by the ATIP-Avenir program, Inserm-Transfert, Inserm, Région Occitanie, University of Montpellier, and Retina France.

## Notes

### Competing Interest Statement

The authors have declared no competing interest.

