## Supplemental Figure legends for "AAV2/9-mediated gene transfer into murine lacrimal gland leads to a long-term targeted tear film modification"

**Supplemental material**

**Fig S1: AAV2/9 and AAV2/5-CAG-GFP transduce LG acinar cells.** Representative images of LG longitudinal sections show GFP protein (green) in acinar cells immunostained for E-Cadh (red) following a single injection of AAV2/9 or AAV2/5-CAG-GFP into murine LG, when compared to AAV2/9 or AAV2/5 empty capsid-injected mice respectively (10^10^ vg / LG in 3 µl, *n* = 3 mice per group). Mice were sacrificed one month post injection. Nuclei are counterstained with Hoechst 33342 (white). Scale bar: 10 µm.

**Fig S2: AAV2/9 and AAV2/5-CAG-GFP transduce LG ductal cells.** Representative images of LG longitudinal sections show GFP protein (green) in ductal cells immunostained for Krt19 (red) following a single injection of AAV2/9 or AAV2/5-CAG-GFP into murine LG, when compared to AAV2/9 or AAV2/5 empty capsid-injected mice respectively (10^10^ vg / LG in 3 µl, *n* = 3 mice per group). Mice were sacrificed one month post injection. Nuclei are counterstained with Hoechst 33342 (white). Scale bar: 20 µm.

**Figure S3**. **mNGF level is increased after a corneal abrasion**. Quantification by ELISA of mNGF level in tears (ng / ml) after a corneal abrasion, and measured 1 day before abrasion, 1, 3 and 7 days post abrasion (*n* = 6 mice). Results are expressed as the mean ± SD. Statistical analysis uses repeated measure one-way ANOVA test followed by Dunnett’s comparisons test. * *p* = 0.0292 and ** *p* = 0.0019 between between 1 day before abrasion and 3 days post abrasion, and between 1 day before abrasion and 7 days post abrasion respectively. ns, not significant. Source data are provided as a Source Data file.

**Figure S4**. **Kinetic study of the secreted mNGF in tears after injection of AAV2/9 and AAV2/5-CAG-mNGF into murine LG**. mNGF protein levels were analyzed by ELISA in the tears of AAV2/9 or AAV2/5-CAG-mNGF-injected mice (10^10^ vg / LG in 3 µl, *n* = 5 mice per group), when compared to the tears of AAV2/9 or AAV2/5 empty capsid-injected mice, at 15, 30, 60 and 90 days post injection. Graph shows the kinetic of mNGF level (ng / ml) in the tears of injected mice. Results are expressed as the mean ± SD. Statistical analysis using repeated measure two-way ANOVA test followed by Tukey’s comparisons test. **** *p* < 0.0001 between AAV2/5 or AAV2/9 empty capsid and AAV2/9-CAG-mNGF groups, *** *p* = 0,0001 between AAV2/9-CAG-mNGF and AAV2/5-CAG-mNGF groups. ns, not significant. Source data are provided as a Source Data file.

**Figure S5. Injection of AAV2/9 empty capsid into murine LG does not alter tear volume nor protein concentration in tears.** AAV2/9 empty capsid were injected in the murine LG (10^11^ vg / LG in 3 µl, *n* = 3). The volume of tears (**A**) and the protein concentration in tears (**B**) were measured 1 day before injection, 7, 30, 60, 120 and 180 days post injection. **A**. Graph showing the volume of tears (µl / min) collected 1 day before injection and at the indicated days post injection. Results are expressed as the mean ± SD. Statistical analysis using Friedman’s one-way ANOVA test followed by Dunn’s multiple comparisons test. ns, not significant. **B**. Graph showing the protein concentration in tears (mg / ml) collected 1 day before injection and at the indicated days post injection. Results are expressed as the mean ± SD. Statistical analysis using Friedman’s one-way ANOVA test followed by Dunn’s multiple comparisons test. ns, not significant.

Source data are provided as a Source Data file.
