## Supplementary figures and images for "AAV2/9-mediated gene transfer into murine lacrimal gland leads to a long-term targeted tear film modification"

### Figures

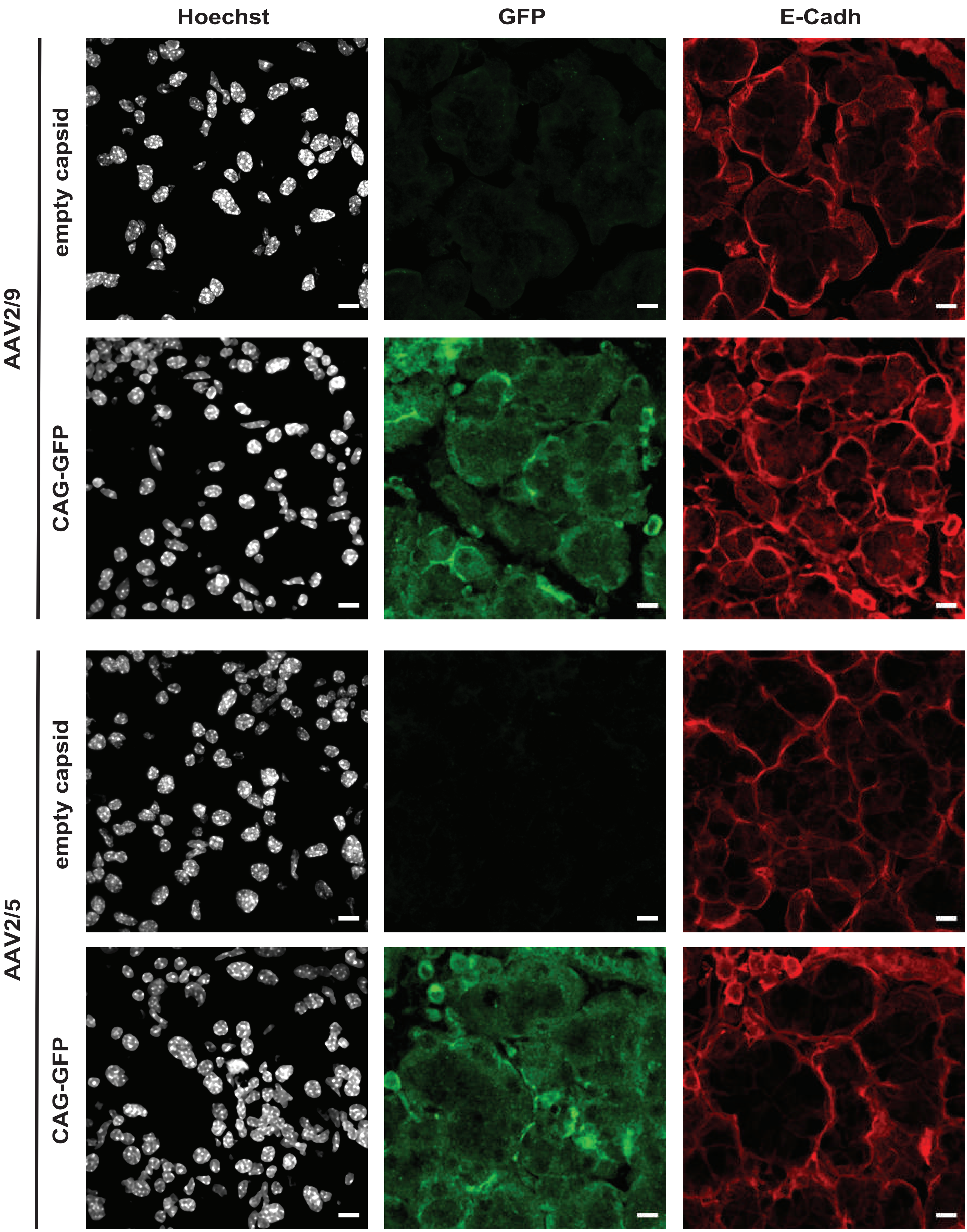

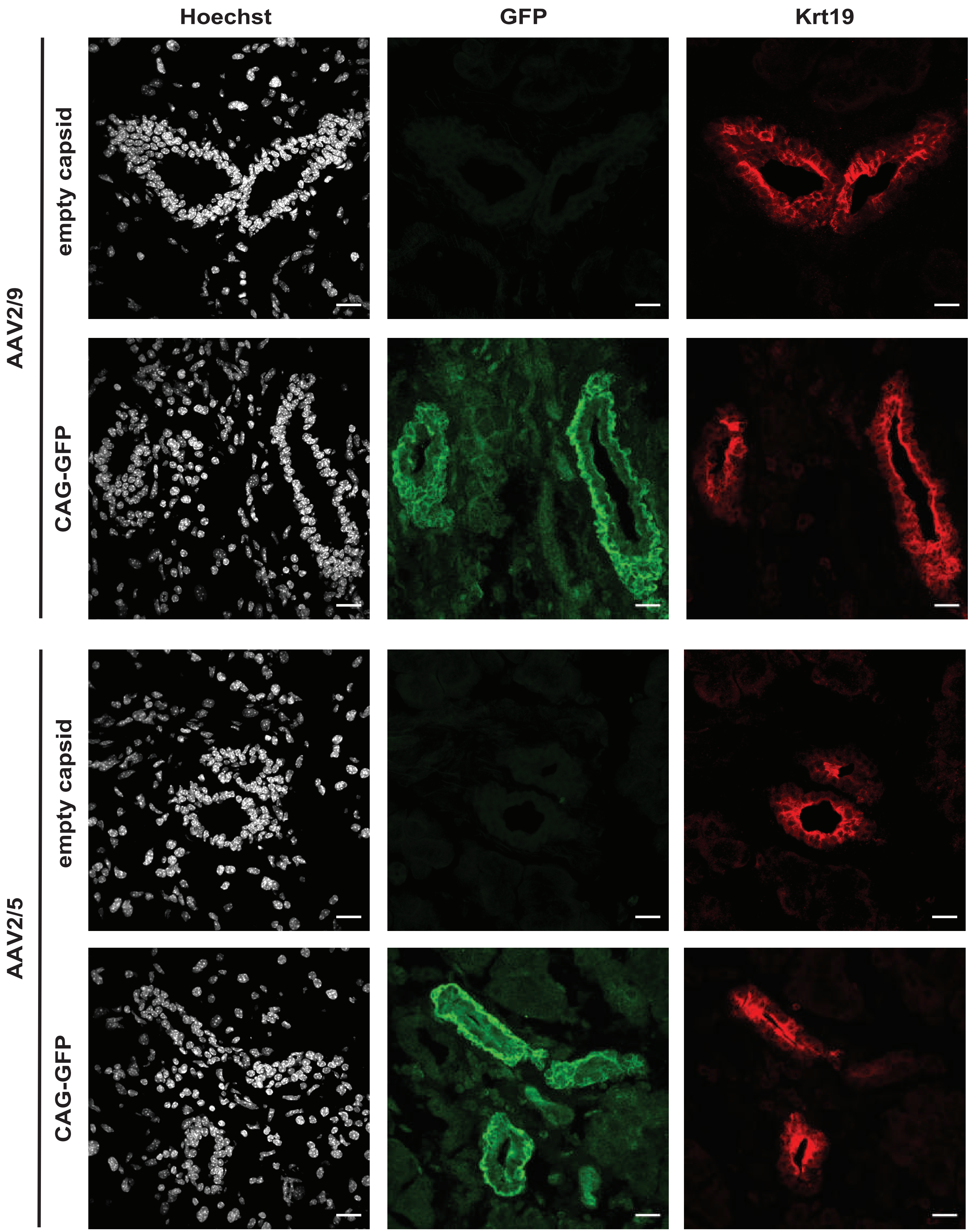

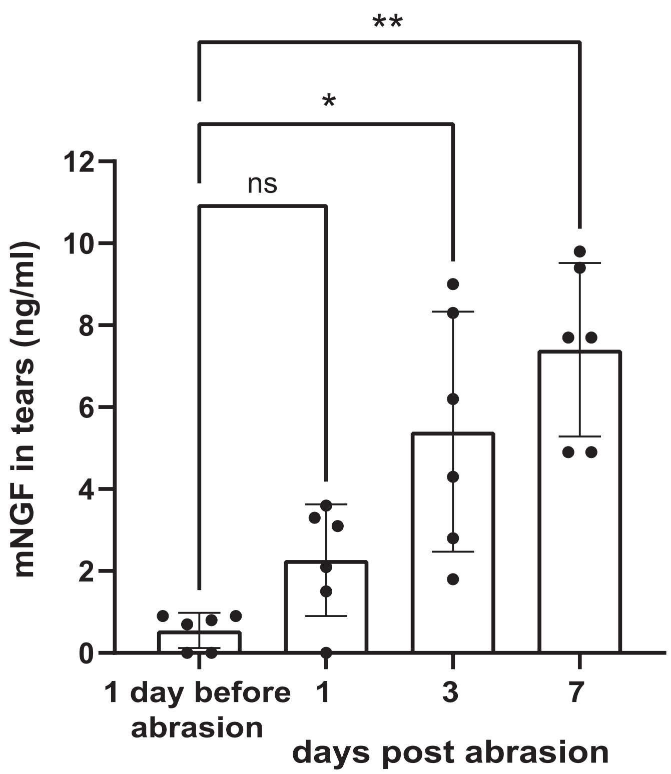

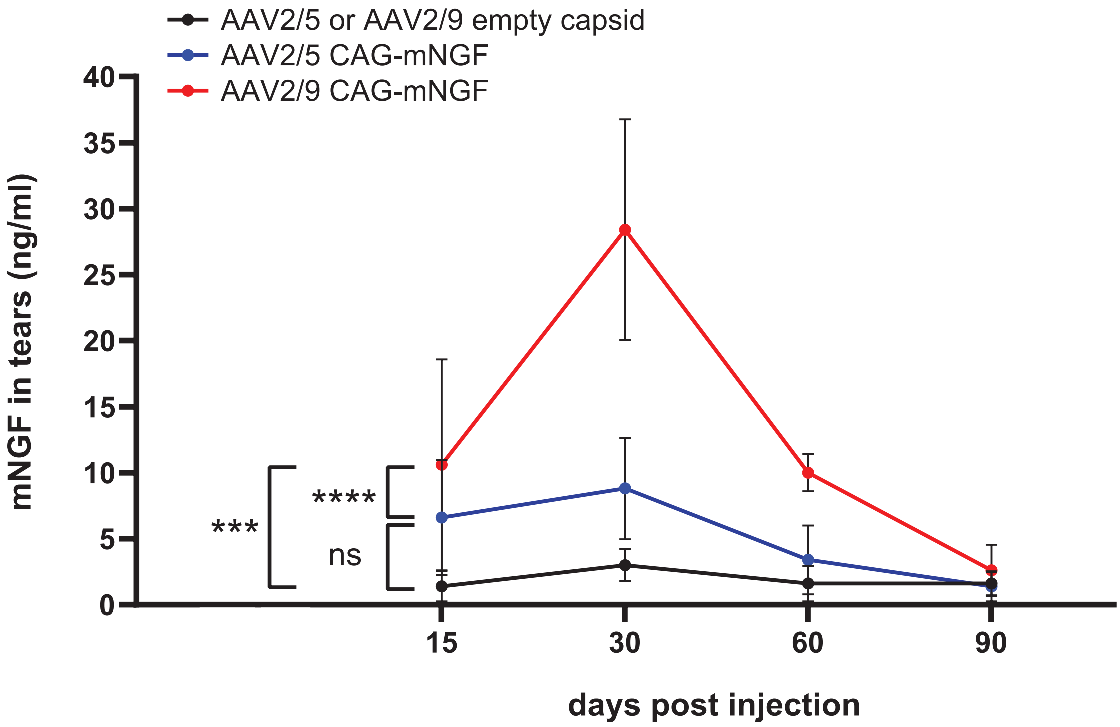

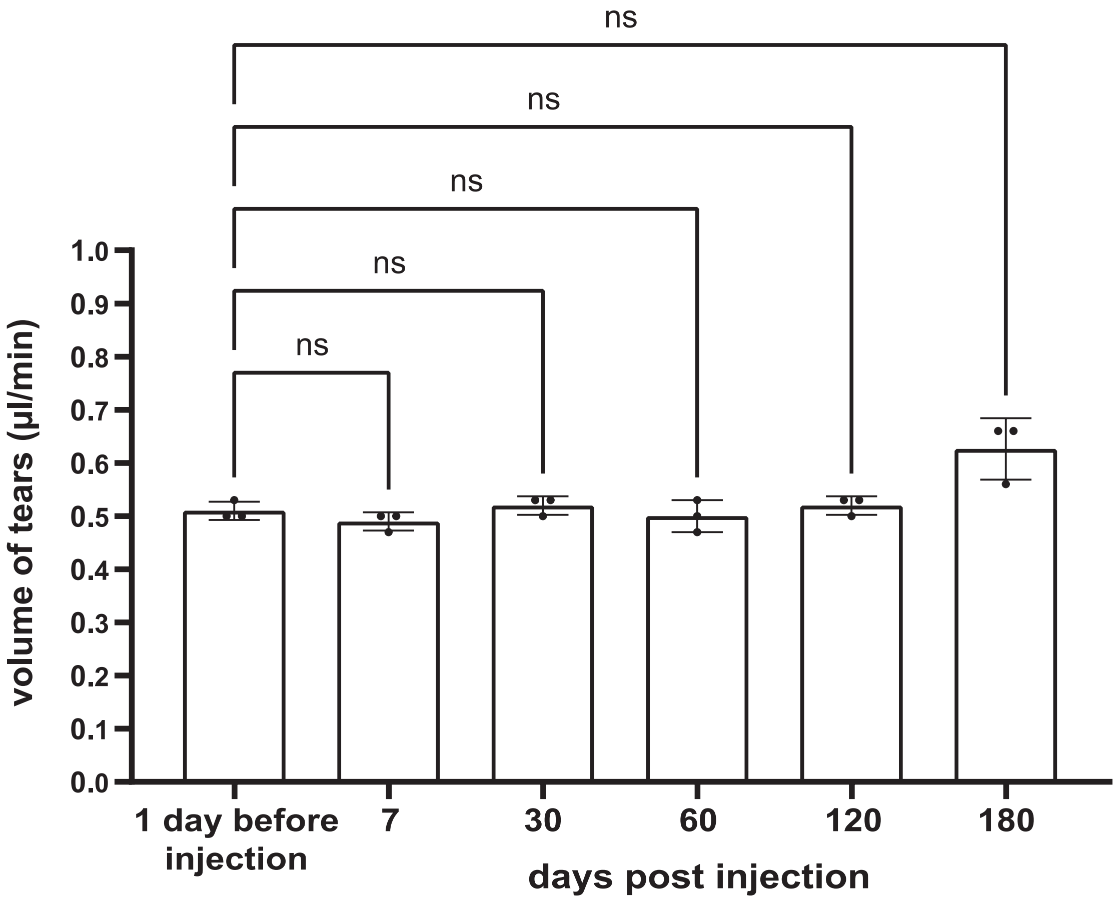

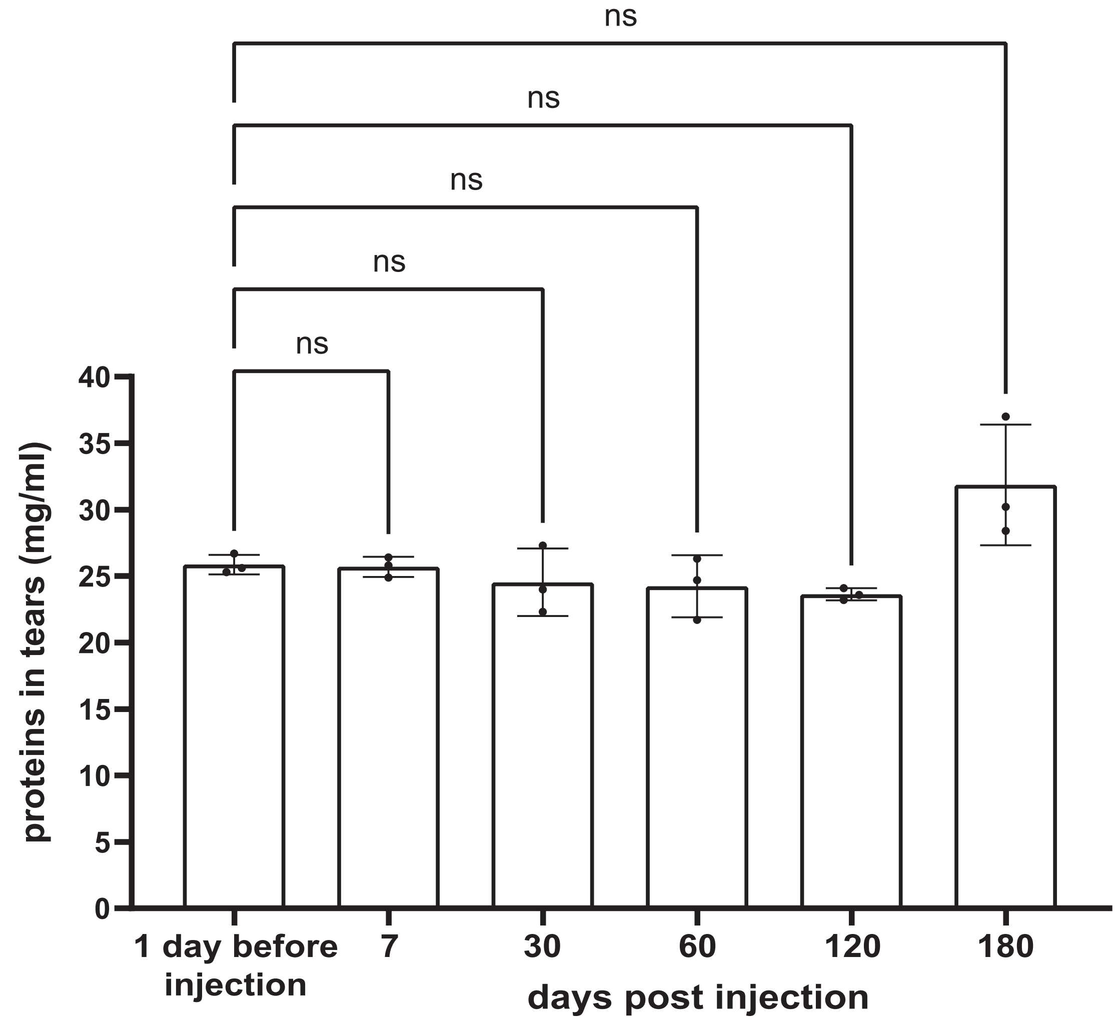
